# The structure and robustness of tripartite ecological networks

**DOI:** 10.1101/2021.10.05.463170

**Authors:** Virginia Domínguez-García, Sonia Kéfi

## Abstract

Until recently, most ecological network analyses have focused on a single interaction type. In nature, however, diverse interactions co-occur, each of them forming a layer of a ‘multilayer’ network. Data including information on multiple interactions has recently started to emerge, giving us the opportunity to have a first glance at possible commonalities in the structure of these networks. We studied the structural features of 44 tripartite ecological networks from the literature, each composed of two layers of interactions (e.g. herbivory, parasitism, pollination), and investigated their fragility to species losses. We found that the way in which the different layers of interactions are connected to each other affect how perturbations spread in ecological communities. Our results highlight the importance of considering multiple interactions simultaneously to better gauge the robustness of ecological communities to species loss and to more reliably identify the species that are important for robustness.

## INTRODUCTION

The ecological network literature has long been dominated by studies of networks containing a single interaction type [de Ruiter et al., 1995, Neutel, 2002, Bascompte et al., 2003]. These studies have contributed considerable understanding about their structure as well as important relationships between the structure and functioning of ecological communities. However, it has become increasingly clear that species in nature are connected by a myriad of interaction types simultaneously, and that considering networks which include the diversity of interaction types could greatly improve our knowledge of the structure and dynamics of ecological communities [Fontaine et al., 2011, Kéfi et al., 2012, Kéfi et al., 2015, Kéfi et al., 2016, García-Callejas et al., 2017, Kéfi, 2020].

The idea of considering multiple types of interactions is not new, and a number of previous studies have investigated the effect of including multiple interaction types on the functioning of ecological communities, especially on their stability [May, 1972, Mougi and Kondoh, 2012, Allesina and Tang, 2012, Sauve et al., 2013, Lurgi et al., 2015, McWilliams et al., 2019, Hale et al., 2020]. Yet, these studies have so far been dominantly theoretical. With the recent publication of the first multi-interaction empirical networks, we begin to know more about their structure [Melián et al., 2009, Pocock et al., 2012, Sauve et al., 2016, Kéfi et al., 2016, Genrich et al., 2016, Dáttilo et al., 2016, Astegiano et al., 2017, Mello et al., 2019], and how it affects persistence [Melián et al., 2009, Kéfi et al., 2016] and robustness [Pocock et al., 2012, Evans et al., 2013, Dáttilo et al., 2016]. In particular, studies on multi-interaction networks have provided new information on whether including several interactions can significantly alter robustness [Dáttilo et al., 2016] and the way perturbations spread through ecological communities [Pocock et al., 2012]. However, in spite of these pioneering studies, there is currently no consensus about the structure of multi-interaction networks or about its consequences for robustness to species loss, in part due to the lack of data-sets, whose amount has only recently started to increase.

Here, we gathered multi-interaction ecological networks currently available in the literature. More specifically, we focused on tripartite networks, i.e. ecological networks containing two types of ecological interactions or *interaction layers*, each of the bipartite kind. Our data-set consists of 44 tripartite networks, each composed by two bipartite networks that include mutualistic (pollination, seed-dispersal and ant-mutualism) and antagonistic (herbivory and parasitism) interactions [Macfadyen et al., 2009, Melián et al., 2009, Pocock et al., 2012, Dáttilo et al., 2016, Hackett et al., 2019, Shinohara et al., 2019]. Using this data-set, we investigated their structural features, identifying possible generalities across interaction types as well as singularities specific to a given interaction type, and we studied the consequences of these structural properties for the robustness of these networks to species loss, focusing on plants, the only group of species common to all our networks and whose disappearance can potentially harm all other species groups.

## METHODS

### Definitions

Tripartite ecological networks are composed of two interaction layers, each of them of the bipartite kind. They therefore contain three different *species sets* (e.g. plants, pollinators and herbivores in a pollination-herbivory network). Among the three species sets, one acts as a link between the two interaction layers (e.g. plant species interact both with pollinators trough pollination and with herbivores trough herbivory). We call this set of nodes that may have interactions in both interaction layers the *linking set*, and the nodes in the linking set with interactions in the two interaction layers the *connector nodes* (see Fig. 1.A and B.).

**Figure 1:**
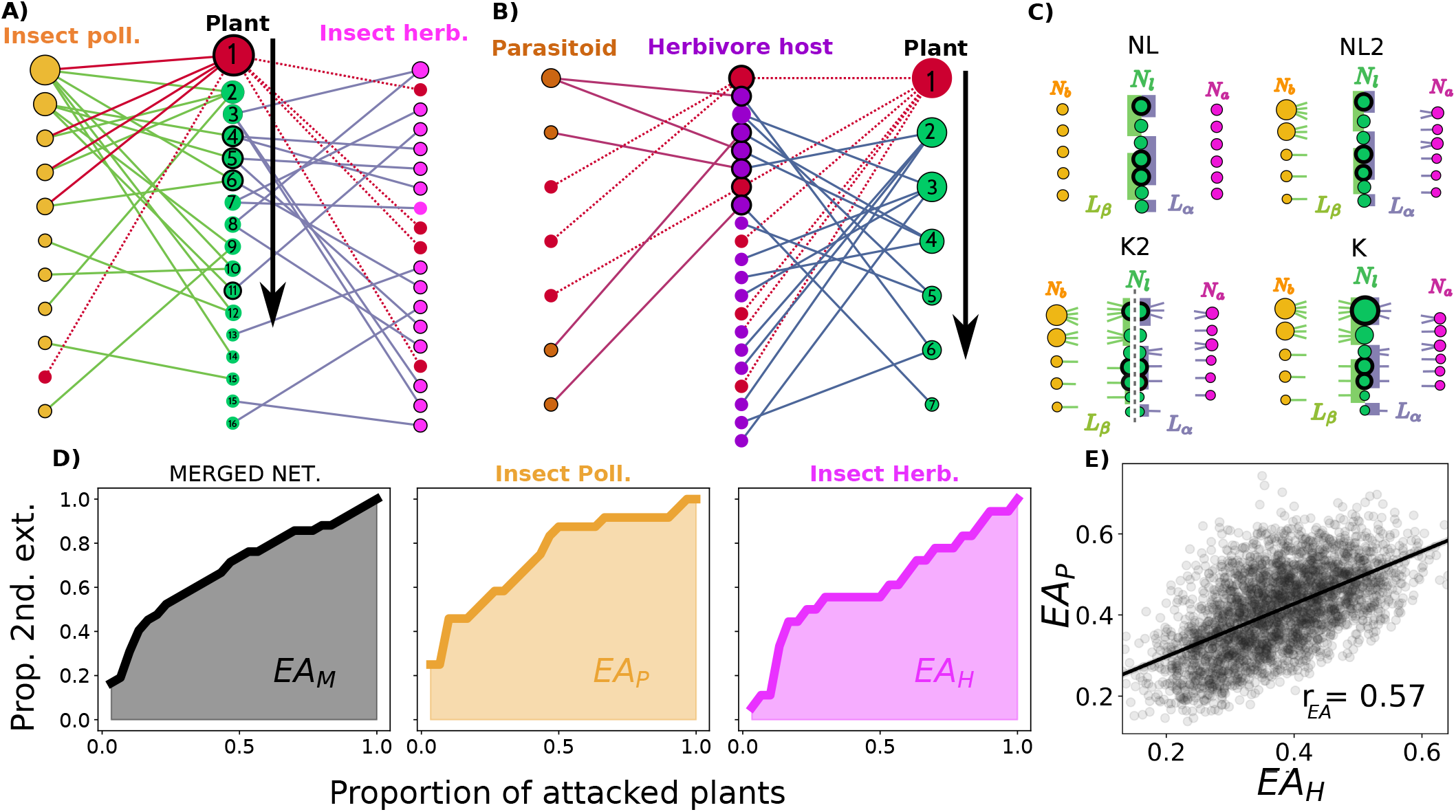
A) Example of a pollination-herbivory tripartite network where plants are the linking set. B) Example of a herbivory-parasitoids tripartite network where herbivores are the linking set. Node colours represents the three different sets of species, and link colours the different interaction layers. Connector nodes in the linking set are highlighted in black. Extinction protocol: plants (green nodes) are progressively removed from the community in the prescribed order, their corresponding links are erased (colored in red) and animal species are declared extinct (colored in red) whenever they lose all their connections. C) The 4 different null models used in this study. Each panel represents what is kept fixed in each null model. D) Extinction curves showing the fraction of extinct animal species in the merged community, in pollinators, and in herbivores respectively, as a function of the number of sequentially removed plants for a given extinction sequence in network A. Different extinction sequences lead to different extinction areas. The larger the area, the more impact a given extinction sequence of plants has on the community. E) Correlation between the extinction areas (*r*_*EA*_) of the two animal species set in the example of network A for 3000 extinction sequences.

### The data-set

We gathered from the literature multipartite ecological networks which included at least two different types of interactions and at least 5 connector nodes. The data-set contains 44 unweighted networks from 6 studies (see references in Table 1). Each network is composed of two ecological bipartite interactions including mutualistic (pollination, seed-dispersal and ant-mutualism) and antagonistic interactions (herbivory and parasitism). We divided the networks in three types according to the signs of the interactions involved: mutualism-mutualism (MM; 3 networks) if both interactions are positive, antagonism-antagonism (AA; 24 networks) if both interactions are negative, and mutualism-antagonism (MA; 17 networks) if one interaction is positive and the other negative (see SI S1).

**Table 1:** Tripartite networks composing the data-set. Columns indicate from left to right: the name of the network (based on the publication it comes from), the sign of the two interactions composing the network (mutualism-mutualism: MM, antagonism-antagonism: AA, mutualism-antagonism: MA), the name of the two interactions layers composing the network (ant-mutualism: A, herbivory: H, parasitoidism: Pa, pollination: P, and seed-dispersal: SD), the d gree heterogeneity 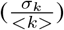, the degree-degree correlations (r), Z-score of degree heterogeneity in the NL null model 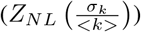, Z-score of degree-degree correlations in K null model (*Z*_*K*_(*r*)), and the reference to the study where the data was gathered.

| name | sign | int | $\frac{\sigma_k}{\langle k \rangle}$ | r | $Z_{NL}(\frac{\sigma_k}{\langle k \rangle})$ | $Z_K(r)$ | Ref. |
| --- | --- | --- | --- | --- | --- | --- | --- |
| Sinohara_1_ALL_PH | MA | H-P | 1.08 | -0.15 | 13.77(**) | 0.73() | [Shinohara et al., 2019] |
| Sinohara_2_ALL_PH | MA | H-P | 1.04 | -0.21 | 12.96(**) | -0.59() | [Shinohara et al., 2019] |
| Sinohara_3_ALL_PH | MA | H-P | 1.21 | -0.25 | 16.53(**) | -0.42() | [Shinohara et al., 2019] |
| Sinohara_4_ALL_PH | MA | H-P | 1.21 | -0.37 | 11.93(**) | -3.31(**) | [Shinohara et al., 2019] |
| Sinohara_ALL_A_PH | MA | H-P | 1.22 | -0.27 | 12.94(**) | -0.97() | [Shinohara et al., 2019] |
| Sinohara_ALL_E_PH | MA | H-P | 1.22 | -0.21 | 21.51(**) | -0.32() | [Shinohara et al., 2019] |
| Sinohara_ALL_I_PH | MA | H-P | 1.26 | -0.24 | 16.21(**) | -0.47() | [Shinohara et al., 2019] |
| Sinohara_2_E_PH | MA | H-P | 0.97 | -0.26 | 7.86(**) | -1.18() | [Shinohara et al., 2019] |
| Sinohara_3_E_PH | MA | H-P | 1.21 | -0.32 | 15.10(**) | -1.27() | [Shinohara et al., 2019] |
| Sinohara_4_I_PH | MA | H-P | 0.84 | -0.17 | 3.51(**) | -0.06() | [Shinohara et al., 2019] |
| Melian_OO_OO_PH | MA | H-P | 2.05 | -0.44 | 72.48(**) | 0.80() | [Melián et al., 2009] |
| Hackett_1_ALL_PH | MA | H-P | 2.16 | -0.39 | 62.36(**) | -0.40() | [Hackett et al., 2019] |
| Hackett_2_ALL_PH | MA | H-P | 1.06 | -0.34 | 9.78(**) | -0.61() | [Hackett et al., 2019] |
| Hackett_1_S_PH | MA | H-P | 1.61 | -0.63 | 10.67(**) | 1.00() | [Hackett et al., 2019] |
| Hackett_1_GL_PH | MA | H-P | 1.55 | -0.46 | 20.63(**) | -0.99() | [Hackett et al., 2019] |
| Pocock_OO_OO_PH | MA | H-P | 1.92 | -0.47 | 67.46(**) | -0.44() | [Pocock et al., 2012] |
| Melian_OO_OO_HSD | MA | H-SD | 1.82 | -0.52 | 59.15(**) | -0.92() | [Melián et al., 2009] |
| McFayden_ALL_A_HP | AA | H-Pa | 1.56 | -0.23 | 75.52(**) | -1.01() | [Macfadyen et al., 2009] |
| McFayden_1_A_HP | AA | H-Pa | 1.31 | -0.38 | 25.98(**) | -1.70() | [Macfadyen et al., 2009] |
| McFayden_2_A_HP | AA | H-Pa | 1.31 | -0.38 | 24.63(**) | -2.79(*) | [Macfadyen et al., 2009] |
| McFayden_3_A_HP | AA | H-Pa | 1.25 | -0.32 | 19.40(**) | -1.77() | [Macfadyen et al., 2009] |
| McFayden_4_A_HP | AA | H-Pa | 1.02 | -0.29 | 13.98(**) | -0.64() | [Macfadyen et al., 2009] |
| McFayden_5_A_HP | AA | H-Pa | 1.12 | -0.27 | 18.96(**) | -1.18() | [Macfadyen et al., 2009] |
| McFayden_6_A_HP | AA | H-Pa | 1.14 | -0.24 | 17.60(**) | -0.39() | [Macfadyen et al., 2009] |
| McFayden_7_A_HP | AA | H-Pa | 1.14 | -0.30 | 20.14(**) | -1.08() | [Macfadyen et al., 2009] |
| McFayden_8_A_HP | AA | H-Pa | 1.23 | -0.31 | 23.94(**) | 0.20() | [Macfadyen et al., 2009] |
| McFayden_9_A_HP | AA | H-Pa | 1.13 | -0.35 | 16.61(**) | -0.92() | [Macfadyen et al., 2009] |
| McFayden_10_A_HP | AA | H-Pa | 1.15 | -0.35 | 15.06(**) | 0.27() | [Macfadyen et al., 2009] |
| McFayden_ALL_B_HP | AA | H-Pa | 1.50 | -0.22 | 68.74(**) | -0.81() | [Macfadyen et al., 2009] |
| McFayden_1_B_HP | AA | H-Pa | 1.29 | -0.28 | 28.94(**) | -0.02() | [Macfadyen et al., 2009] |
| McFayden_2_B_HP | AA | H-Pa | 0.98 | -0.22 | 11.12(**) | -0.15() | [Macfadyen et al., 2009] |
| McFayden_3_B_HP | AA | H-Pa | 1.29 | -0.26 | 24.19(**) | -0.95() | [Macfadyen et al., 2009] |
| McFayden_4_B_HP | AA | H-Pa | 1.12 | -0.37 | 16.11(**) | -1.80() | [Macfadyen et al., 2009] |
| McFayden_5_B_HP | AA | H-Pa | 1.19 | -0.25 | 18.72(**) | -0.36() | [Macfadyen et al., 2009] |
| McFayden_6_B_HP | AA | H-Pa | 0.84 | -0.17 | 11.41(**) | 0.11() | [Macfadyen et al., 2009] |
| McFayden_7_B_HP | AA | H-Pa | 0.86 | -0.23 | 12.07(**) | -0.85() | [Macfadyen et al., 2009] |
| McFayden_8_B_HP | AA | H-Pa | 1.11 | -0.38 | 14.43(**) | -1.28() | [Macfadyen et al., 2009] |
| McFayden_9_B_HP | AA | H-Pa | 1.00 | -0.30 | 11.07(**) | -0.78() | [Macfadyen et al., 2009] |
| McFayden_10_B_HP | AA | H-Pa | 1.22 | -0.34 | 21.62(**) | -0.78() | [Macfadyen et al., 2009] |
| Hackett_1_ALL_HP | AA | H-Pa | 0.81 | -0.31 | 6.43(**) | 0.20() | [Hackett et al., 2019] |
| Hackett_1_WL_HP | AA | H-Pa | 0.66 | -0.46 | 0.43() | -0.65() | [Hackett et al., 2019] |
| Melian_OO_OO_PSD | MM | P-SD | 1.87 | -0.35 | 48.77(**) | -3.20(**) | [Melián et al., 2009] |
| Dattilo_OO_OO_PSD | MM | P-SD | 1.25 | -0.29 | 32.15(**) | -3.16(**) | [Dáttilo et al., 2016] |
| Dattilo_OO_OO_PA | MM | P-A | 1.26 | -0.22 | 33.96(**) | -2.41(*) | [Dáttilo et al., 2016] |

### Null models and Z-scores

To assess the importance of network structure in determining a certain network feature, we compared measurements of that feature performed on empirical networks with measurements performed on randomized versions of those networks keeping some properties fixed. We used four different null-models (Fig. 1.C), which – going from the least to the most constraining – are as follows: “NL” keeps the number of species constant in each species set and the number of links constant in each interaction layer, “NL2” adds the constraint of keeping the degree distribution of the animal nodes constant, “K2” keeps the degree distribution of animals and plants but not the total degree of the linking set species (i.e. it breaks the correlation between the degree of the linking set speciesin the two interaction layers), and “K” keeps the degree of each node constant while links are reshuffled within a layer (see SI S2).

To compare the value of a given metric in an empirical network with that obtained in the random ensemble, we measured the Z-Score of that metric. Z-Scores quantify the number of standard deviations by which the value of a raw score (i.e., the value measured in the empirical network) is above or below the mean value measured in the random ensemble, and are defined as:

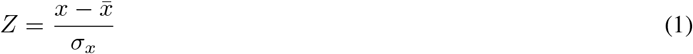

where 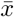 is the average value of x in the random ensemble and *σ*_*x*_ its standard deviation. If the Z-Score is between −1.96 and +1.96 (*p* > 0.05) one cannot reject the null hypothesis with a 95% confidence level; For a measure to be outside the 98% confidence interval (*p*<0.02) the absolute value of the Z-score has to be larger than 2.33.

### Basic structural features: Degree distribution and correlations

Since the data-set is very heterogeneous, we measured the heterogeneity of the degree distribution 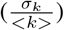. If this value is close to or higher than one, it means that the variance in the degree is of similar order as the average degree [Bell, 1992]. We compared the degree heterogeneity of our empirical networks with that obtained in the random ensemble of 200 networks built using null model “NL” by measuringthe Z-score of the degree heterogeneity.

We next look at degree-degree correlations. In many real networks, nodes are not connected randomly, but instead there is a significant bias for nodes of high degree to either connect to nodes of high degree (‘assortative’ networks) or to nodes of low degree (‘disassortative’ networks) [Newman, 2003]. This tendency is usually measured using Pearson’s correlation coefficient (*r*):

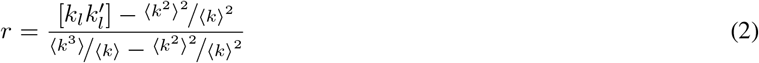

where ⟨ · ⟩ represent averages over all nodes of the network, and [*k*_*l*_*k*_*l*_′] is the product of the degree of each of the two nodes (*k*_*l*_, *k*_*l*_′) belonging to link *l* over all links in the network. To test whether empirical networks are more or less assortative/disassortative than expected by chance given their degree distribution, we measure the Z-Score of Pearson’s correlation coefficient with respect to the values obtained in the random ensemble of 200 networks built using null model “K”.

### Structural metrics of the connector nodes

We were interested in studying how the two different interactions of the tripartite networks are interconnected trough the connector nodes. We used three metrics to quantify this:

- The proportion of connectors nodes in the linking set (*C*), i.e. the proportion of species in the linking set that have links simultaneously in the two interaction layers [Astegiano et al., 2017].
- The participation ratio (*PR*_*C*_). This species-level metric quantifies whether the links of node *i* are primarily concentrated in one interaction layer or if they are well distributed among the two[Battiston et al., 2014]. We quantified the PR as two times the ratio between the lowest degree in both interaction layers divided by the total degree of the node 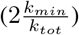. Hence *PR* = 1 if the links are perfectly split among the two interaction layers, and it approaches 0 as the split grows more uneven. We obtained the PR of the connector nodes by computing the average value of their PR.
- The proportion of connector hubs (*H*_*C*_), i.e. the proportion of linking set hubs, meaning the 20% of the species in the linking set with the highest degree, that are connector nodes (i.e. that have links in both interaction layers).

### Robustness of tripartite networks

We investigated how the loss of plant species affected the tripartite networks in our data-set. As in previous studies [Pocock et al., 2012, Dáttilo et al., 2016], we focused on plants because they are the only group of species whose extinction can possibly affect all other species groups. To quantify fragility to plant loss, we followed the protocol shown in Fig.1A and B: we sequentially removed plants in a given order (the ‘extinction sequence’) keeping track of the number of secondary extinctions of animal species at each step. We considered that an animal species undergoes extinction when it has lost all its links. It is worth noting that secondary extinctions work differently in mutualistic-antagonistic and mutualistic-mutualistic networks compared to antagonistic-antagonistic networks. In the former, after removing a plant, all herbivores that no longer have resources go extinct and so do all pollinators that don’t have any resources left, which means that erasing a plant may generate *simultaneous* secondary extinctions in the two animal species set (Fig. 1.A). In antagonistic-antagonistic networks, herbivores are linking set species, so when a plant disappears, all herbivores without resources go extinct, which may subsequently trigger extinctions of parasitoids. In this case, removal of a plant will generate *cascading* extinctions(Fig. 1.B).

By plotting the proportion of extinct animal species as a function of the proportion of deleted plant species and measuring the area under the curve, we obtained the ‘extinction area’, a number between 0 and 1 (Fig. 1.D). The extinction area is a standard way of measuring the efficiency with which a given extinction sequence tears down an ecological community [Allesina and Pascual, 2009, Domínguez-García and Muñoz, 2015]: the larger the extinction area, the most impact a given extinction sequence has on the community. Note that robustness is defined as 1– extinction area.

When working with multipartite networks, several extinction areas can be measured depending on the species set on which secondary extinctions are measured [Pocock et al., 2012, Dáttilo et al., 2016]. Here, we measured:

- the extinction area of the tripartite network (*EA*_*M*_): we kept track of the proportion of extinct animal species as a function of the proportion of deleted plant species, where the proportions are measured with respect to the total number of animals (irrespective of their species set) and plants.
- the extinction areas of the two animal species sets (*EA*_*P*_, *EA*_*H*_): we measured the proportion of extinct animal nodes with respect to the total number of animals in each species set (e.g. how many pollinators undergo extinction compared to the original number of pollinators), and the proportion of deleted plants is measured with respect to the total number of plants in the tripartite network.
- the extinction areas of the two bipartite networks. In this case, the tripartite network is split in two bipartite networks, on which the same protocol as above is performed. These two networks are not identical to the two interaction layers because the linking set species that are not connected in a given layer are not considered in the bipartite network, which affects the calculation of the extinction area. We thereby obtain two extinction areas (*EA*_*L*_ and *EA*_*S*_, respectively for the smaller and larger networks, in terms of species number). Note that in AA networks, the protocol can be performed only on the herbivory network since there is no direct connection between plants and parasitoids.

We applied 3000 random extinction sequences of plants to each of the tripartite networks in the data-set, and for each extinction sequence we measured the different extinction areas indicated before. Here, results are presented for random extinction sequences but EA for other extinction scenarios (increasing or decreasing degree of plants) are presented in SI S6.

This protocol gives a distribution of values rather than a number, so we use the Z-test to compare the distribution of the extinction area in the empirical and random ensembles of networks. We compute the Z-score as:

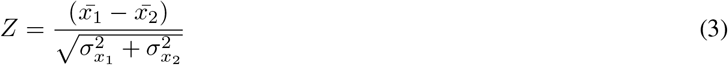

where 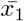 is the mean value of sample one (empirical), 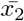 is the mean value of sample two (randomized ensemble), 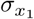 is the standard deviation of sample one divided by the square root of the number of data points, and 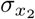 is the standard deviation of sample two divided by the square root of the number of data points.

### Interdependence

We measured the correlation between the extinction areas of the two different species sets (other than plants) in the tripartite networks [Pocock et al., 2012], hereafter called ‘interdepence’ (*r*_*EA*_) (Fig.1.E). When we drive plants to extinction, a ‘high’ correlation between the extinction areas of pollinators and herbivores implies that the same plants that are relevant for one of the species set (i.e. those that generate a larger extinction area in pollinators) will also be relevant in the other species set (i.e. will also generate a large extinction area in herbivores). If, on the other hand, the relevant plants are not the same in the two species sets, we expect a low correlation between extinction areas.

### Plant importance rankings

The importance of each plant species for the different animal sets and for the whole community (i.e. tripartite network) was quantified based on the correlation coefficient between the extinction area when plants were removed in random order and the position of the plant in the extinction sequence [Pocock et al., 2012]. The rationale is that the ‘importance’ of a plant cannot be directly assessed from the number of secondary extinctions caused by its loss because if lost at the start (rather than at the end) of the extinction sequence, fewer secondary extinctions are expected; however, if a plant is ‘important’, then the extinction area is expected to be higher when it is lost earlier in the sequence than when it is lost later. Hence, the higher the extinction area caused by any extinction sequence, the better that extinction sequence actually resembles the importance of plants for the survival of the community. To obtain the plant importance rankings (three in total: one for each of the two interaction layers and one for the whole community), we ranked each plant species by increasing correlation between its order of appearance in extinction sequences and the corresponding extinction areas (i.e. plants that have a larger negative correlation are considered more important; Fig. 4A-G).

To asses to what extent one of the two interaction layers drives the robustness of the whole community, we measured the similarity between the importance of a plant in the pollination (respectively herbivory) layer and the importance of a plant in the whole community, namely *S*_*P*_ (*S*_*H*_). We quantified *S*_*int*_ as the square of the correlation coefficient between the ranking of the plant in each interaction layer and the ranking in the whole community.

We then classified the networks in three categories: those where one layer was driving the process (one *S*_*int*_ was above 0.9, meaning that 90% of the variance in the importance ranking in the whole community can be traced to one of the two rankings), those where the ranking in the whole community was a mixture of the rankings in both interaction layers (both *S*_*int*_ were between 0.5 and 0.9) and finally those where the importance ranking was emergent (both *S*_*int*_ were below 0.5, meaning that no interaction layer ranking was able to explain at least 50% of the ranking of importance in the whole community).

### Estimating the merged extinction area

We also tested whether one can express the EA of the whole community (*EA*_*M*_) as a combination of the extinction area of the two independent bipartite network composing them. To do that we performed the following linear regression:

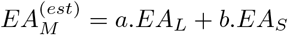

where *EA*_*L*_ and *EA*_*S*_ are respectively the extinction areas of the two bipartite networks composing the tripartite network under study, the larger (i.e. with more species) and the smaller one.

### Multiple regressions

We performed a multiple regression of interdependence and extinction area based on the structural features of the tripartite networks using the package *statsmodel* in Python. We selected the structural features that were more relevant for the interdependence or the extinction area by choosing the model with a lowest AIC.

## RESULTS

### A few generalist and many specialist species

The study of basic structural features, namely degree heterogeneity and degree-degree correlations, revealed that the empirical tripartite networks share two characteristics: they have heterogeneous degree distributions both at the species set and at the whole network (‘aggregated’) level, and their degree-degree correlations are negative, both at the interaction layer level and at the aggregated level.

Indeed, pulling all species together (i.e. independent of their species set), all empirical tripartite networks in the database show a higher degree heterogeneity than expected in the null model “NL” (Table 1). This feature, caused by the presence of generalist and specialist species (SI S3, Fig. S5), remains true in the different species sets, although there are differences among them, with parasitoids showing the lowest degree heterogeneity, and herbivores and plants the highest.

We also found the empirical tripartite networks to be disassortative, i.e. they show negative degree-degree correlations (*r*) : high degree nodes tend to be connected to low-degree nodes and vice-versa (Table 1). This value is consistent with those of the “K” ensemble in the majority (86%) of networks, which suggests that the disassortativity can be traced back to the degree heterogeneity for most empirical networks except for the Mutualistic-Mutualistic (MM) ones. Looking at each of the interaction layers that compose our tripartite networks reveals a similar result (SI S3).

### Connecting the interaction layers

To investigate how the two interaction layers connect to form the tripartite networks, we focused on three structural features of the linking set nodes (Methods), based on which we found fundamental differences between the different types of tripartite networks (Fig. 2).

**Figure 2:**
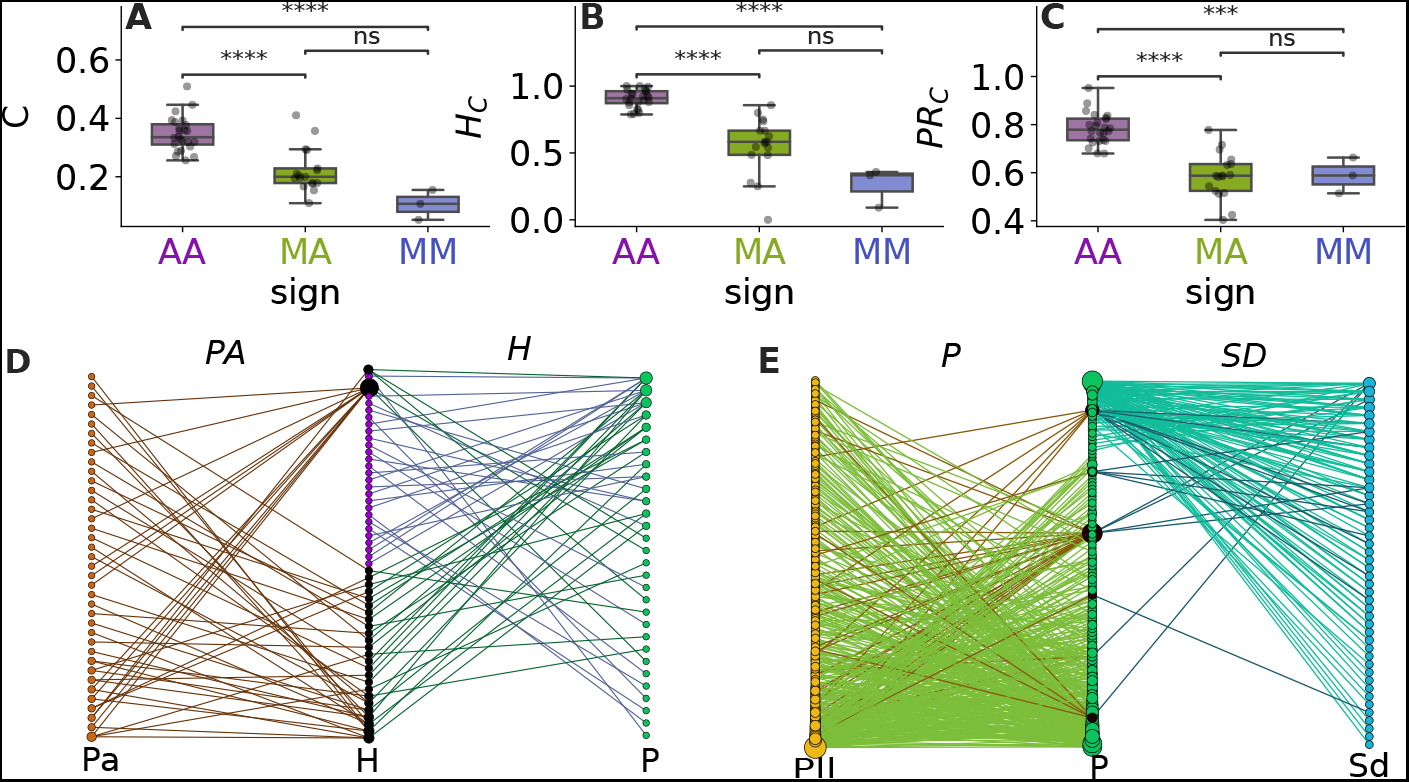
A) Proportion of connector nodes in the linking set, B) Proportion of linking set hubs that are connector nodes, C) Average PR of the connector nodes. Boxplots are color-coded by network type: Antagonistic-Antagonistic: AA, Mutualistic-Antagonistic: MA and Mutualistic-Mutualistic: MM. Differences among categories are measured by independent t-tests (**** p<0.001, *** p<0.01, *ns* not significant). D-E: Two contrasting examples of empirical networks in our data-set. D) An AA network of Herbivory (H)-Parasitoidism (Pa). E) A MM network of Pollination (P)-Seed-dispersal (SD).

In Antagonistic-Antagonistic (AA) networks, ~35% of the linking set species (herbivore hosts) are involved in both parasitic and herbivory interactions (i.e. are connector nodes). Moreover, on average, 96% of the linking set hubs are connectors, and they have their links equally split among the two interaction layers (average PR of 0.89).

We found a very different pattern in mutualistic-mutualistic (MM) networks (Fig. 2 A-C) for which only ~10% of the linking set species (plants) are involved simultaneously in the two types of mutualistic interactions, and only 32% of linking set hubs act as connector nodes. Also, the connector nodes have their links less equally split among the two interaction layers (average PR of 0.59).

Mutualistic-Antagonistic (MA) networks are not significantly different from MM networks and tend to have values of the structural features studied that are intermediate between those of AA and MM networks (Fig. 2 A-C). About ~22% of the linking set species are involved simultaneously in the two types of mutualistic interactions, ~ 56% of linking set hubs act as connector nodes and the average PR is ~ 0.59.

### Interdependence of the robustness of animal species

Following recent studies on the robustness of multipartite networks [Pocock et al., 2012, Evans et al., 2013], we investigated whether the robustness of different species sets in the tripartite networks were correlated, i.e. if they were interdependent (Methods). We found that, in general, when plants were driven to extinction in a random order, interdependence (*r*_*EA*_) was either positive or null (Fig. 3.A), with fundamental differences between AA networks and the other two types of networks. The value of interdependence found in AA networks was on average significantly higher from that found in MM and MA networks, which is consistent with our results on hubs and connectors, suggesting that the two layers of these networks are strongly interconnected. However, this correlation is not significantly higher from what is expected in the null models (SI S5). Since this correlation is already present in all the null models investigated, regardless the type of constraint this suggests that this positive correlation is due to the particular layout of these networks, more specifically, to the cascading extinction process.

**Figure 3:**
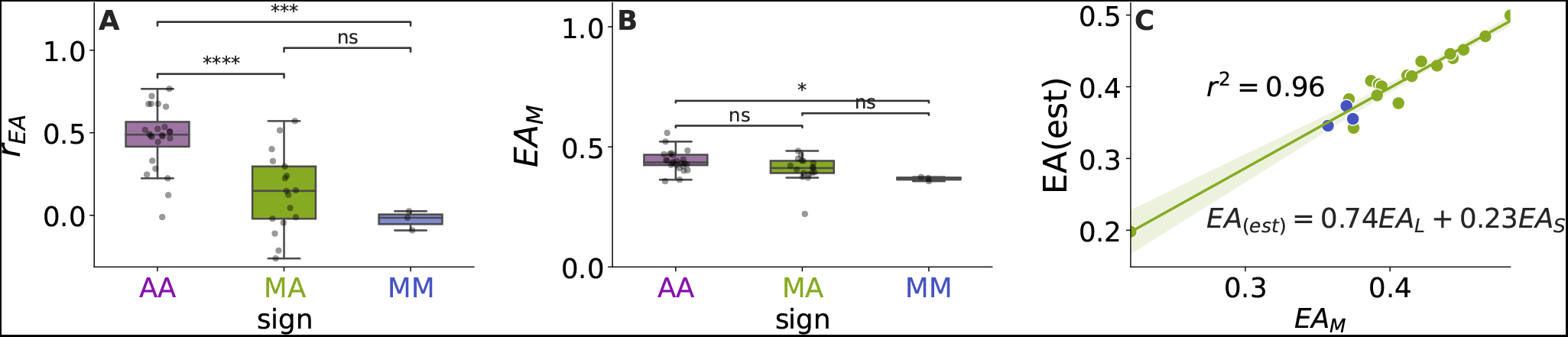
A: Boxplot of interdependence in the different types of tripartite networks. B: Extinction area of the tripartite networks in our data-set (*EA*_*M*_) when plants are randomly driven to extinction. All boxplots are color-coded based on the type of tripartite network (AA, MA or MM). Differences among the categories are measured by independent t-tests (**** p<0.001, *** p<0.01, *ns* not significant). C: Estimated merged extinction area (*EA*_(*est*)_) vs merged extinction area (*EA*_*M*_) in the empirical MA and MM networks of our database. The text is the best estimation of the merged extinction area as a combination of the extinction areas of the larger (*EA*_*L*_) and smaller (*EA*_*S*_) bipartite networks that compose the tripartite network.

**Figure 4:**
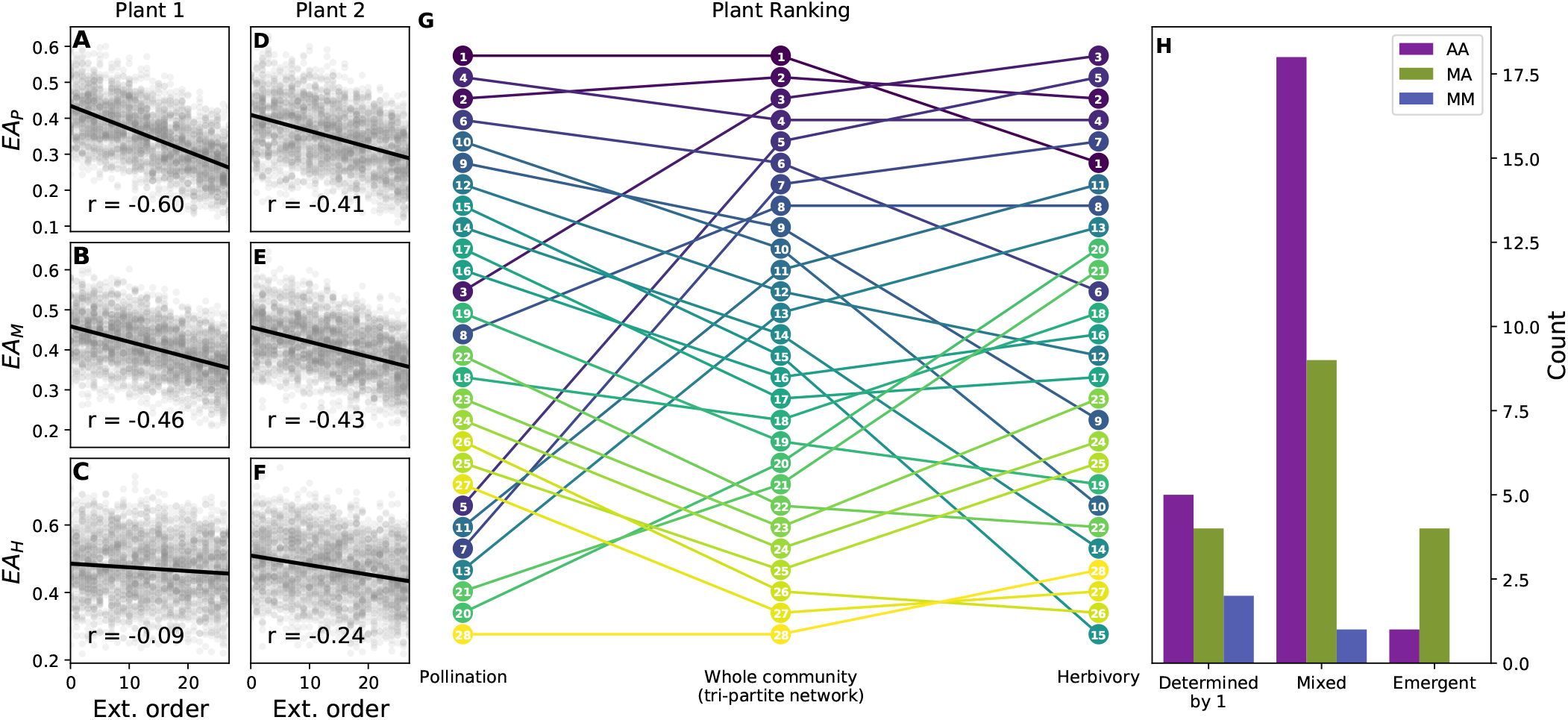
A-F Scatter plot of the Extinction area in the pollination layer (*EA*_*P*_), merged tripartite network (*EA*_*M*_), and herbivory layer (*EA*_*H*_) vs the order of two plants (plant 1 and plant 2) in the extinction sequence. The correlation coefficients are used to determine the ranking of importance of plant species. G: Ranking of plant importance in the herbivore layer (left), whole community (center) and pollination layer (right). Each plant is represented by a disk with a number which is their order in the ranking of importance of the whole community (in the tripartite network). The height of the disk represents its order in each of the three different rankings (i.e. the higher the position, the more important). Lines between balls are a help to track changes in the rankings. H: Classification of the tripartite networks in our database according to *S*_*int*_ (similarity between the ranking of plant importance in the whole community and in the interaction layers), attending at if the ranking of plant importance in the whole community is mainly determined by only one interaction layer, it is a mixture of the rankings of importance in the two interaction layers (mixed), or if does not resembles any of the rankings of importance in the interaction layers (emergent).

In MM and MA networks, the interdependence is close to null, meaning that the extinction areas of the two layers seem largely decoupled from each other. Comparison with the null models suggests that, in this case, the degree heterogeneity of the observed networks makes them more interdependent than expected (since null models that don’t control for degree heterogeneity, namely “NL” and “NL2”, predict negative interdependence).

Studying how interdependence relates to the three structural features studied revealed differences among network types as well. More specifically, in AA networks, interdependence is correlated (albeit weakly) with the proportion of connectors (*C*), while in MA and MM it varies with the proportion of connector hubs (*H*_*C*_) and their (un)balanced involvement in the two interaction layers (*PR*_*C*_) (Table 2).

**Table 2:** Table of regression of structural features on interdependence (*r*_*EA*_) and extinction area (*EA*)

| | $r_{EA}$ | | $EA$ | |
| --- | --- | --- | --- | --- |
|  | AA(best) | MA & MM(best) | AA(best) | MA & MM(best) |
| $\sigma_k / \langle k \rangle$ | | | -0.68***<br>(0.14) | -0.38*<br>(0.19) |
| $r_b$ | | -0.30**<br>(0.14) | | |
| $C$ | 0.40*<br>(0.20) | | -0.25**<br>(0.11) | 0.51**<br>(0.19) |
| $H_C$ | | 0.58***<br>(0.14) | | |
| $PR_C$ | | 0.34**<br>(0.13) | 0.25*<br>(0.14) | |
| AIC | 67.92 | 38.84 | 41.61 | 53.24 |
| Observations | 24 | 20 | 24 | 20 |
| $R^2$ | 0.16 | 0.73 | 0.76 | 0.38 |
| Adjusted $R^2$ | 0.12 | 0.68 | 0.73 | 0.31 |
| Residual Std. Error | 0.96 | 0.58 | 0.53 | 0.86 |
| F Statistic | 4.20* | 14.15*** | 21.39*** | 5.18** |

Note:
\* $p < 0.1$ ; \*\* $p < 0.05$ ; \*\*\* $p < 0.01$

### Robustness of the tripartite networks

The robustness of AA networks was found to be higher than that of MM networks when plants were randomly driven to extinction (Fig.3.B) although differences among the three types of networks are overall very small. Surprisingly, this suggests that even if the different ways in which the tripartite networks are connected seem to have a significant effect on interdependence, this difference does not translate into significant differences in the global robustness of tripartite networks. In other terms, a higher interdependence between the interaction layers did not cause a higher overall extinction area.

We also found that the extinction area of the whole tripartite network (i.e. the merged extinction area) could be predicted by the extinction areas of the two bipartite networks composing it. The estimated extinction area, a combination of the extinction areas of the two bipartite networks, was in very good agreement (*R*^2^ = 0.96) with the merged extinction area (Fig. 3.C). When the extinction area of only one interaction layer is used, *R*^2^ is at most 0.8 (SI 6, Fig.S10).

The structural features which best determined the merged extinction area were the degree heterogeneity and the proportion of connector nodes in MA and MM networks, and also the PR in AA networks (Table 2).

### Plant importance for robustness

We investigated which plants were more important for the survival of the ecological community and to what extent those plants were the same in the two interaction layers. We built three rankings of plant importance – one for each interaction layer and one for the whole community – in which a plant is considered to be more important if the extinction area is larger when that plant is attacked earlier in the extinction sequence (Methods). For example, a plant can be considered important based on the pollination and whole community rankings (plant 1, Fig. 4.A, B), but not so based on the herbivory ranking (Fig.4.C). Other plants are considered important based on the three rankings (plant 2, Fig.4.D-F).

We studied to what extent the importance of a given plant at the whole community level was driven by its importance based on the interaction layers composing the network (Fig.4.H). In the majority of networks ( ~63%), the importance of a plant in the whole community is a mixture between its importance in the two different interaction layers (i.e. *S*_*int*_ is between 0.5 and 0.9; Methods), while in ~25% of the networks it was mostly driven by the ranking of one of the interaction layers (i.e *S*_*int*_ of one interaction layer is above 0.9). This is especially relevant in MM networks, where 2 out of the 3 networks lie in this category, probably because of the high dissimilarity between the sizes of the two interaction layers (180 pollinators vs 27 seed-dispersers and 173 pollinators vs 30 ants). In a few cases ( ~12%), the ranking of importance in the whole community did not resemble any of the interaction layer’s rankings (i.e. both *S*_*int*_ were below 0.5), meaning that the importance of a plant when the two interactions are considered simultaneously changes dramatically compared to its importance when the interactions are considered separately.

## DISCUSSION

We studied the structure of tripartite ecological networks containing two types of ecological interactions, both to investigate how interactions are connected in different types of tripartite networks, and to study their robustness to loss of plant species. While studies on multi-interaction networks have been gradually appearing in the literature in the last years, this is to our knowledge, the first study comparing several multi-interaction networks, allowing us to reveal commonalitiesas well as particularities.

The study of their basic structural features showed that – consistently with previous analyses on networks with a single interaction [Jordano et al., 2002, Memmott et al., 2004, Solé and Montoya, 2001, Dunne et al., 2002] – all the tripartite ecological networks in our database have a heterogeneous degree pattern, with a few generalist and many specialist species. The heterogeneity in species degrees induced disassortativity, where high degree species tend to be linked to low degree species and vice-versa, a result also consistent with previous studies on single layer ecological networks [Jonhson et al., 2013].

Despite these similarities in the basic structural features, we found fundamental differences in the way the two interaction layers are connected. First Antagonistic-Antagonistic (AA) networks have significantly more connectors than Mutualistic-Mutualistic (MM) and Mutualistic-Antagonistic (MA) networks. Moreover, in AA networks, hubs are almost all connectors (i.e. generalist herbivores tend to have parasitoids), while in MM networks most hubs are not connectors (i.e. generalist plants tend not to be involved in two types of mutualism simultaneously). Intuitively, we expected these differences in the connectivity patterns to affect how perturbations spread through tripartite networks and to eventually impact the robustness of entire networks.

Looking at how the robustness of the two interaction layers of a given tripartite network correlate to each other shows that AA networks tend to show a positive interdependence, while MM and MA networks show a lower interdependence (close to null), especially MM networks. In AA networks, cascading extinctions introduce a positive correlation between the extinction areas. As for MM and MA networks, the fact that the plant hubs are involved in the two interaction layers seems to determine their interdependence (as suggested by the fact that interdependence is significantly correlated with *H*_*C*_ – the proportion of linking set hubs that are connectors – and *PR*_*C*_ – the participation ratio of the connector nodes). These differences in interdependence are expected to have consequences for the way disturbances spread through tripartite networks. These results suggest that in more interdependent communities (i.e. with higher values of *r*_*EA*_) such as those driven by antagonistic interactions, affecting one of the plant species is expected to not only impact herbivores, but also the species that depend on them (e.g. parasites in our data-set). In mutualistic communities, interventions aimed at restoring or protecting a plant species are expected to not automatically benefit animals that are not involved in the same type of mutualism as the targeted plant. Our results add to previous evidence showing that the benefits of an intervention are not always expected to propagate throughout the whole network [Pocock et al., 2012] and highlight the relevance of the type of interactions present in the ecological community to plan restoration efforts. For example, in MM and MA networks, positive cascading effects are expected only if the plant hubs act as connector nodes and are the focus of the restoration plan.

These results raise the question of how these patterns of interconnections between the layers of a network affect its overall robustness. Surprisingly, we found that more interdependent communities were not necessarily more fragile to plant losses. Rather, the overall robustness is a result of the particular structure of the interactions in that community; more specifically, the robustness of MA and MM tripartite networks was found to be a combination of the robustness of the two bipartite networks composing them, stressing the relevance of knowing the structure of connections in both interaction layers to better quantify the robustness of the whole tripartite network. Interestingly, looking at the two interaction layers simultaneously did not result in a dramatic change in the robustness of the whole community, as already reported for one of the networks in the database [Dáttilo et al., 2016]. This contrasts with what was found in dynamical models of ecological communities, where introducing multiple types of interactions can change their behaviour (e.g. coexistence, productivity, stability) drastically [Melián et al., 2009, Allesina and Tang, 2012, Mougi and Kondoh, 2012, Sauve et al., 2016, Miele et al., 2019]. Nonetheless, considering the two interactions simultaneously improves the quantification of the overall robustness and is crucial to identify the most important plants. In most tripartite networks, the ranking of plant importance in the whole community was determined by the importance of plants in both interaction layers (with the exception of MM, that where mostly driven by one interaction layer, pollination, in one pollination-dispersion network and in the pollination-ant mutualism networks, probably because of their disproportionate size and disconnection among interaction layers). In a few cases, considering the whole community could even alter the picture considerably, since the ranking of plant importance in the whole community was emergent, i.e. it was not highly correlated with the ranking of importance in neither of the interaction layers separately.

The results we present here advance our knowledge of how different interactions connect ecological communities, and how that affects the robustness of tripartite networks to plant loss. Taken together our results highlight the need to consider multiple interactions simultaneously to accurately measure the robustness of ecological communities as well as to identify the plant species that are key for robustness. In particular, these results can be of interest when considering strategies for the protection or restoration of ecological communities. In line with previous studies [Pocock et al., 2012], our results suggest that the optimistic view in which protecting one key species could propagate and benefit the whole community does not seem to be justified in MA and MM networks. In particular, the variability exhibited in MA networks suggests that a good knowledge of the interactions taking place in the ecological community is needed to predict whether a positive effect will cascade through the network, as well as to decide which plants should be the focus of restoration efforts.

## Conclusion

Taken together our results suggest that considering multiple ecological interactions simultaneously does not have a dramatic impact on the overall robustness of multi-interaction networks to plant losses. However, a multi-interaction approach is crucial to know the interdependence of the different network layers, to better gauge their robustness, and to correctly determine the importance of the plants at the whole community level. These results advance our knowledge of how multiple bipartite interactions are connected in ecological communities and of how these interconnections affect the spread of perturbations in communities with multiple interaction types.

## Supporting information

Supplementary information to main text

## ACKNOWLEDGEMENTS

This work was supported by the grant ANR-18-CE02-0010-01 of the French National Research Agency ANR (project EcoNet).

