## Supplementary information to main text for "The structure and robustness of tripartite ecological networks"

### SUPPLEMENTARY INFORMATION FOR: STRUCTURE AND ROBUSTNESS OF TRIPARTITE ECOLOGICAL NETWORKS

A PREPRINT

**Virginia Domínguez-García**
ISEM, CNRS, Univ. Montpellier, IRD, EPHE, Montpellier, France


**Sonia Kéfi**
ISEM, CNRS, Univ. Montpellier, IRD, EPHE, Montpellier, France;
Santa Fe Institute, 1399 Hyde Park Road, Santa Fe, NM 87501, USA


October 5, 2021

#### Supporting information Text

#### 1 Data-set compilation

In order to build the data-set of tripartite networks, we gathered networks from 6 different studies (see Fig. S0) containing networks with two or more different types of interactions. When a study included networks from many sites and habitats, we considered all the individual networks as well as the merged versions (e.g. in the Macfayden study [10], we considered the networks of the different organic or traditional farms, and also the two networks merging all the traditional and all the organic farms). When a study contained networks with more than two interactions, we considered all the tripartite networks (composed by only two different interactions) that could be generated combining all the interaction layers in the original study. From all the rendered networks we only considered those with at least 5 species in the linking set, resulting in the 44 networks that form our data set (see Table S1 and Table 1 in the main text). The goal was to obtain as many different networks as possible from each reference, circumventing the problem of having too few linking nodes to carry out statistical calculations. Note that the results shown in the manuscript hold also when removing networks that may contain redundant information. The name of the 44 networks in our data set includes first the name of the leading author of the original study (e.g. McFayden) followed by the number of the site or '00' if there was only one site in the original study, a code describing the habitat type (e.g 'A' or 'B' for the two different types of farms in the study) or '00' if the original study only included one habitat type, and a code describing the two interactions included in the network (e.g. PH for pollination-herbivory, SDH for seed dispersal-herbivory, HPa for herbivory-parasitism, PSD for pollination-seed dispersal and PA for pollination-ant mutualism) (see tables S1 and Table 1).

Table S1: Multipartite networks included in our analyses, indicating the sign of the interactions (i.e. if the tripartite network has both mutualistic and antagonistic interactions (MA), only antagonistic interactions (AA), or only mutualistic interactions (MM)), the two ecological interactions composing the tripartite network, the number of network of each type, and the reference.

| Sign | Interactions (Acronym) | Number of networks | references |
|---|---|---|---|
| MA | herbivory-pollination (H-P) | 16 | [11] [14] [13] [7] |
| | herbivory-seed dispersal (H-SD) | 1 | [11] |
| AA | herbivory-parasitism (H-Pa) | 24 | [10] [7] |
| MM | pollination-seed dispersal (P-SD) | 2 | [4] [11] |
| | pollination-ant mutualism (P-A) | 1 | [4] |

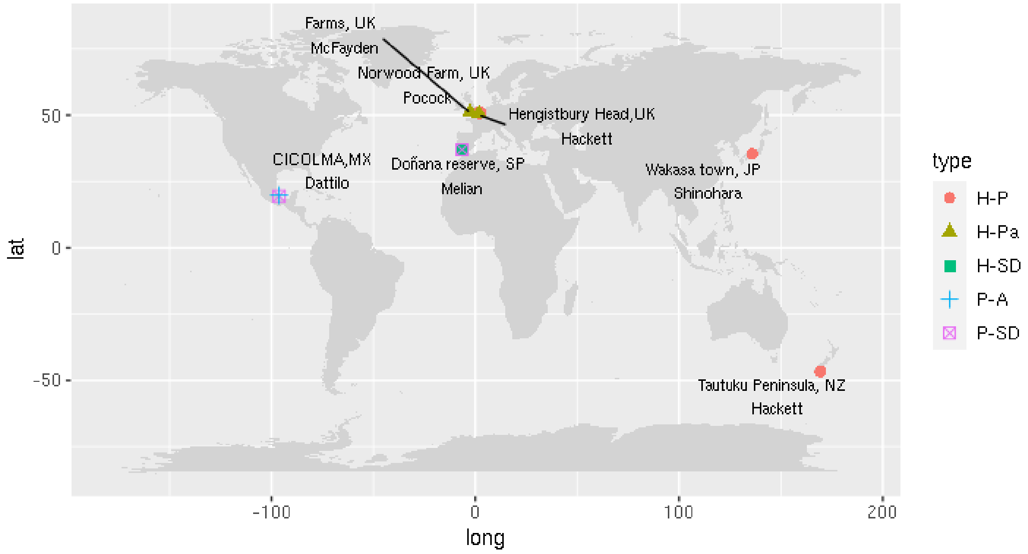


Figure S0: Map showing the location of the 6 studies we used to build the database of networks. The color and shape of the points represents the different types of networks, and the text the name of the site, the country, and the surname of the leading author of the study.

#### 2 Null models and Z-scores

To assess the importance of network structure in determining a certain network feature, we compared measurements of that feature performed on empirical networks with measurements performed on randomized versions of those networks keeping some properties fixed. We use four different null-models (see Fig. S1), going from the less restrictive to the more restrictive (left to right in the figure), they are:

- "Constant NL": the number of nodes in each of the three species sets ($N_a$, $N_b$, and $N_l$), as well as the number of links in each interaction layer ($L_\alpha$, $L_\beta$) are kept constant. The randomization is performed in both interaction layers ($\alpha$ and $\beta$) separately, by linking $L_\alpha(L_\beta)$ pairs of nodes from the two different sets involved in the interaction: a and l (b and l), ensuring that each of the nodes in the original network receives at least one link. Networks generated in the "constant NL" ensemble have the same size and connectance as the empirical networks, but erase any other structure present in the original interactions. In particular, the degree distribution of each species set is lots as well as the degree-degree correlations between nodes of species sets. When a metric is similar in the empirical networks and in the constant NL ensemble, one knows that the number of species and the connectance are the relevant features determining that metric.
- "Constant NL 2": the number of nodes in each species set ($N_a$,$N_b$, $N_l$) is kept constant, as well as the number of interactions in each interaction layer ($L_\alpha$,$L_\beta$), and the degree of all nodes except those in the linking set. This null model allows to study the effect of the degree distribution of the linking set species has in a given metric. When a metric is similar in the empirical network and in this random ensemble, one knows that the size of the sets and connectance are the the relevant features determining that metric (i.e. the degree distribution of the linking set species does not play a relevant role).
- "Constant K 2": the number of nodes in each species set ($N_a$,$N_b$, $N_l$) is kept constant, as well as the degree distribution of the species in both interaction layers. However, the nodes in the linking set do not keep their total degree (i.e. we beak the correlation between the degree of the linking set nodes in each interaction layer). This null model allows to study the effect the correlation between the degree of connector nodes in each interaction layer has in a given metric. When a metric is similar in the empirical network and in this ensemble one knows that the correlation in the degrees of the connector species in different interaction layers is not relevant in determining that given metric.
- "Constant K": the number of nodes in each species set ($N_a$,$N_b$, $N_l$), the total number of links in each interaction layer ($L_\alpha$,$L_\beta$), and each node's degree (the number of links of a given node) are kept constant. However, the identity of neighbours are reshuffled within a layer. The reshuffling of the empirical networks is performed using the 'curveball algorithm' [17]. Networks generated in the "constant K" ensemble have exactly the same degree distribution as the empirical networks, but erase any other structure present in the original interactions, in particular the degree-degree correlation. When a metric is significantly different in the empirical network and in this null

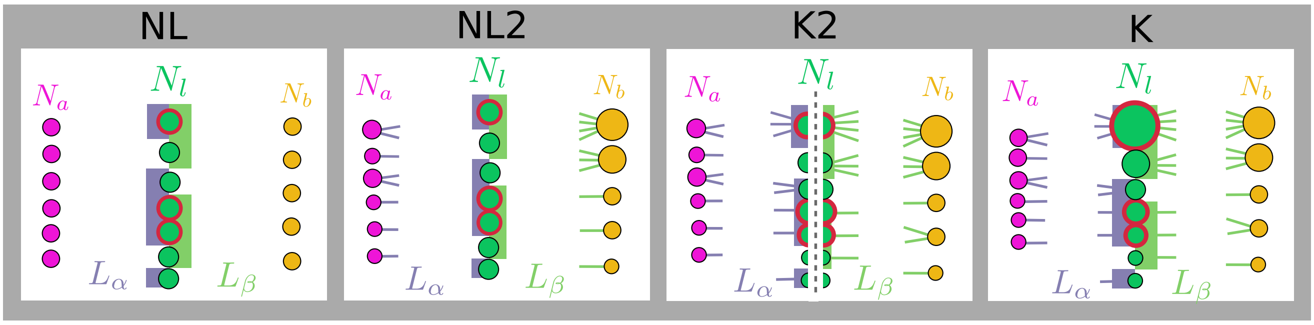


Figure S1: The 4 different null models used in this study. Each figure represents what is kept fixed in each null model, going from the less restrictive on the left, to the more restrictive on the right. The color of the nodes represent the different species set, the colour of the link the two different ecological interactions, the size of the node is proportional to its degree (when kept), and connector nodes are highlighted in red.

model, one knows that it is due to other structural features than degree distribution. When a metric is similar in the empirical networks and in the constant K ensemble, one knows that size and degree distribution are the relevant features determining that metric.

To compare the value of a given metric in an empirical network with that obtained in the random ensemble, we measure the Z-Score of that metric. The Z-Score quantifies the number of standard deviations by which the value of a raw score (i.e., the value measured in the empirical network) is above or below the mean value of what is being measured in the random ensemble, and is defined as:

$$Z = \frac{x - \bar{x}}{\sigma_x} \tag{1}$$

where $\bar{x}$ is the average x measured in the random ensemble and $\sigma_x$ its standard deviation. Positive/negative values mean that the measure in the empirical network is higher/lower than in the random-ensemble. The relevant Z score values when using a 95% confidence level are -1.96 and +1.96. The p-value associated with a 95% confidence level is 0.05. If the Z-Score is between -1.96 and +1.96, the p-value will be larger than 0.05, and one cannot reject the null hypothesis; Conversely, for a measure to be outside the 98% confidence interval (p<0.02) the absolute value of the Z-score has to go up to 2.33.

In the case where one wants to compare two distributions, for example when comparing the extinction area in the empirical network and the random ensemble (note that the extinction area of the empirical networks is a distribution of values rather than a number) we use the z-test. In this case, we compute the Z-score as:

$$Z = \frac{(\bar{x_1} - \bar{x_2})}{\sqrt{\sigma_{x_1}^2 + \sigma_{x_2}^2}} \tag{2}$$

where $\bar{x_1}$ is the mean value of sample one (empirical network), $\bar{x_2}$ is the mean value of sample two (randomized ensemble), $\sigma_{x_1}$ is the standard deviation of sample one divided by the square root of the number of data points, and $\sigma_{x_2}$ is the standard deviation of sample two divided by the square root of the number of data points. .

#### 3 Heterogeneous degree distributions and disassortativity

We measure basic structural features (degree heterogeneity and degree-degree correlations) in the tripartite networks in our data set. We find that they have a highly heterogeneous degree distribution, with an average value of $\sigma_k/<k>$ of 1.26 (see Fig. S2.A). Grouping networks according to the sign of the two ecological interactions involved, we find small differences among them, with MA networks being slightly more heterogeneous than AA networks (see Fig. S2.A), but the differences are not significant. Comparing the degree heterogeneity of the networks to that measured in the "constant NL" random ensemble (where the number of nodes in each species set and the number of links in each interaction layer are kept constant) shows that all the observed networks have a degree distribution significantly more heterogeneous than the null expectation, with Z-Score values over 1.96 in all networks but one, independent of the interaction type (see Fig.S2.C below and Table 1 in the main text).

This is also true when looking at the average degree heterogeneity by species set (see Fig. S3.A and C) or by interaction layer (Table S3). This high degree heterogeneity means that empirical tripartite networks, contrary to the expectation in random networks where all nodes have a similar connectivity, have few nodes with a high connectivity (generalists) and many nodes with a lower connectivity (specialists), as shown in Fig. S4. This 'long tailed' degree distribution is a well established feature of plant-pollinator [9, 12], plant-seed disperser [9] and, to a lesser extent, food webs [15, 5] that we recover in the tripartite networks in our data-set as well. Finally, looking only at the degree heterogeneity of the nodes in the linking set, we find that in 50% of networks the degree heterogeneity of the nodes in linking set in the tripartite network is higher than the degree heterogeneity of the subset of linking set nodes in the two bipartite networks (Table S2), and in all cases the degree heterogeneity of the linking set nodes in the tripartite is at least larger than that of one of the two subsets (Table S2). This means that the two layers are connected in such a way that the degree heterogeneity of the linking set nodes is increased.

For more detailed information on the Z-scores of the degree heterogeneity by species set and by interaction layer for each network, see Tables S6 and S7 at the end of the Appendix.

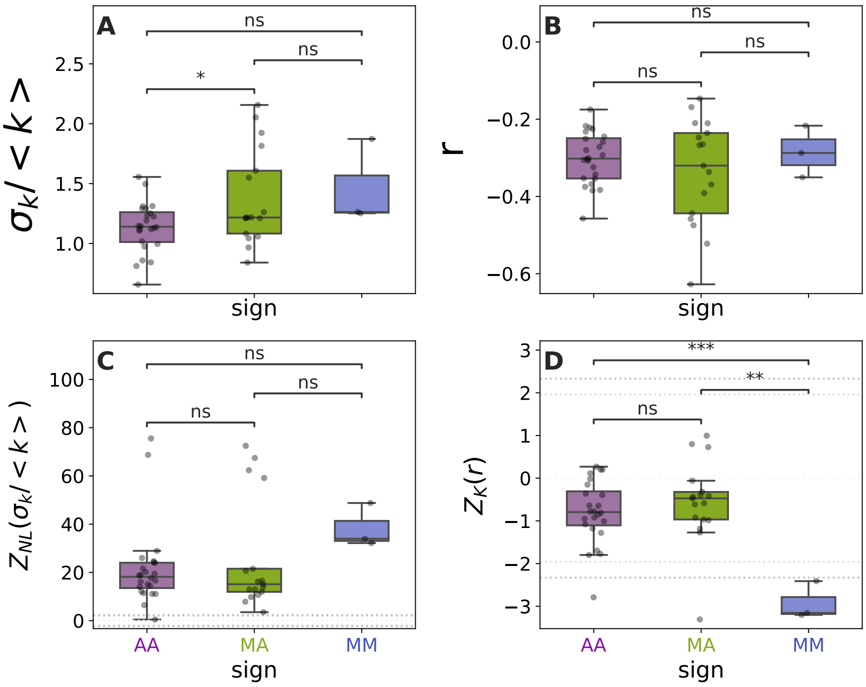


Figure S2: Boxplots of the basic structural features in them empirical tripartite networks of our data-set grouped (and color coded) by the sign of the interactions involved. A: Degree Heterogeneity ($\sigma_k/ < k >$) B: Degree-degree correlations ($r$). C: Z-score of the degree heterogeneity ($\sigma_k/ < k >$) in the "constant NL" null model. D: Z-score of the degree-degrees correlations ($r$) in the "constant K" null model in the tripartite networks. In B and D horizontal grey lines mark the limits of the confidence interval of 1.96 (95%) and 2.33 (98%). Differences between groups are measured by independent t-test (*** $p < 0.001$, ** $p < 0.01$, 'ns' not significant).

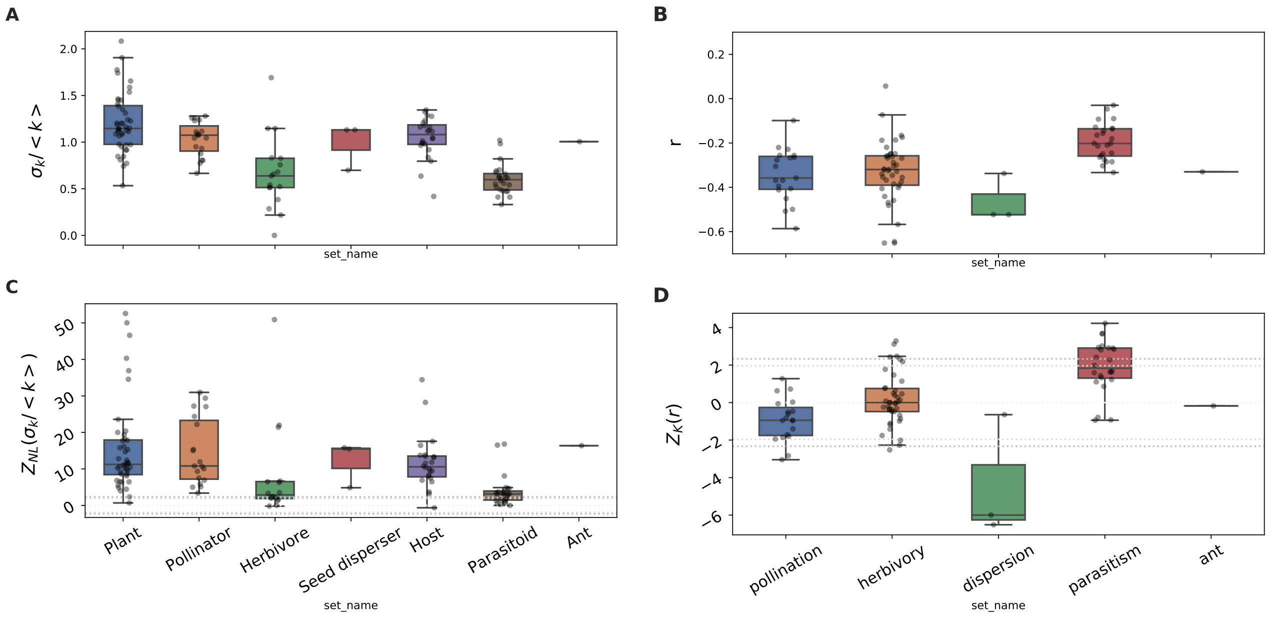


Figure S3: A: Degree heterogeneity of the empirical tripartite ecological networks ($\sigma_k / < k >$) at the species set level B: Degree-degree correlations ($r$) at the interaction layer scale. C: Boxplot of the Z-score of the degree heterogeneity ($\sigma_k / < k >$) by species set in the "constant NL" null model. D: Boxplot of the Z-score of the degree-degree correlations by interaction layer in the "constant K" null model. Horizontal grey lines mark the limits of the confidence interval of 1.96 (95%) and 2.33 (98%) for rejecting the null hypothesis.

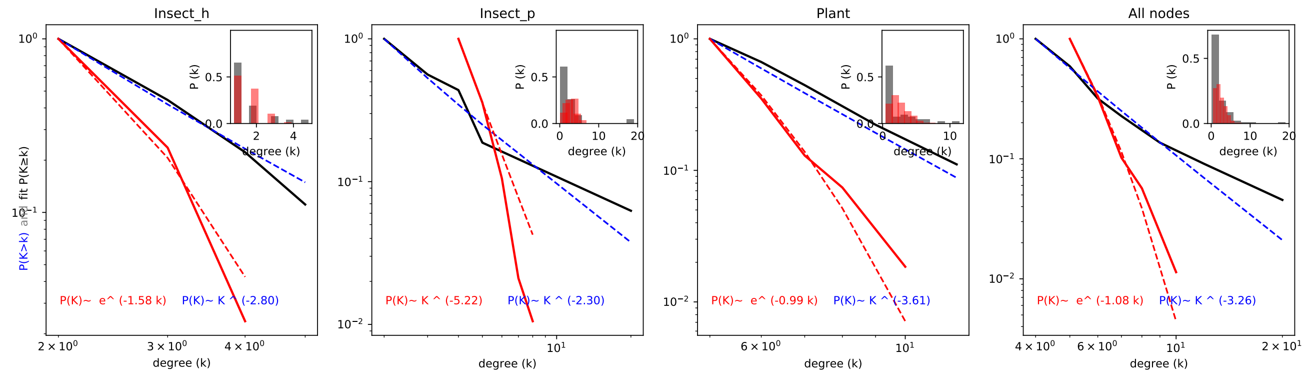


Figure S4: Cumulative degree distribution in a multipartite 'mutualistic-antagonistic' network of pollination-herbivory compared with its randomization in the 'constant NL' null model. Figures show the degree distribution for the three different sets of species, from left to right: insect herbivores, insect pollinators, plants and all nodes combined (i.e. the merged network). The continuous black line is the empirical cumulative degree distribution, with the best power law distribution fit in a dashed blue line. The continuous red line is the cumulative degree distribution in the "constant NL" ensemble (100 randomizations), and the dashed red line the best exponential fit to that distribution. The insets show the degree distribution of the empirical network (in grey) and that of 100 randomizations (in red).

Apart from this heterogeneity in the species degree, in empirical networks the species are not randomly connected, rather the degree of a node can depend on the degree of its neighbours. This is known as 'mixing patterns' or degree-degree correlations, and is measured using correlation coefficients between the degree of a node and the degree of its neighbours. Pearson's correlation coefficient between the degree of a node and the average degree of the node's neighbours ($r$) was found to be negative in all empirical networks, indicating that they are all disassortative, i.e. that species with many connections tend to be linked to species with few connections (see Fig. S2.B). Since degree heterogeneity is known to promote degree disassortativity [8], we also studied if the empirical networks were more disassortative than expected given their degree distribution by comparing the observed values to those of the "constant K" ensemble. In line with previous findings in ecological plant-mutualistic networks [8], we find that most of the observed tripartite networks are not significantly more disassortative than the null expectation (with the exception of MM networks, which exhibit a higher disassortativity than expected), as the value of $r(k)$ in the empirical networks is consistent with those of the "constant K" ensemble average with a confidence con 95%(98%) in 86%(90%) of the networks (see Fig. S2.D above and Table 1 in the main text).

When we look at the interaction layer scale (i.e. at the two unilayer networks composing the tripartite network), we find that all pollination, dispersion, ant-mutualism and a majority (80%) of herbivory networks are disassortative, while most of parasitoids networks (80%) are assortative (i.e. have $r > 0$), although most herbivory and parasitism networks exhibit weak degree-degree correlations (values of $r$ close to 0). However, as with the tripartite networks, when we compare with the randomizations in the 'constant K' null ensemble they tend to not be significantly more assorative/disassortative than the null expectation (Table S3), with 75%(91%) or interaction layers having a value of $r(k)$ compatible with that of the random ensemble with a 95%(98%) confidence interval. This means that most of the assortativity/disassortativity found in the empirical networks stems from their low/high degree heterogeneity. Nevertheless, looking with more detail at the different interacting layers shows that the parasitism interaction layers tend to be rather more assortative than the null expectation, while dispersion and pollination tend to be more disassortative than the random expectation, and we find both trends in herbivory, with herbivory layer from MA networks showing a tendency to being slightly more assortative than the null expectation, and the contrary in herbivory layers from AA networks. Finally, by comparing the degree-degree correlation of the two interaction layers composing the tripartite network and the degree-degree correlations of the merged tripartite networks we see that in all networks but one the merged network is more disassortative than one of the two interaction layers that compose it, and that in 55% of networks (almost all the AA networks, one MA and one MM) the combination of the two interaction layers is made such that the degree-degree correlations of the tripartite networks are more negative (i.e. more disassortative).

| | name | name_layer_A | name_layer_B | LS_HD | lsA_HD | lsB_HD | Increased LS HD |
|---|---|---|---|---|---|---|---|
| 0 | Sinohara_1_ALL_PH | herbivory | pollination | 0.95 | 1.01 | 0.77 | |
| 1 | Sinohara_2_ALL_PH | herbivory | pollination | 0.97 | 0.63 | 0.93 | * |
| 2 | Sinohara_3_ALL_PH | herbivory | pollination | 1.15 | 1.00 | 0.80 | * |
| 3 | Sinohara_4_ALL_PH | herbivory | pollination | 1.20 | 1.05 | 1.04 | * |
| 4 | Sinohara_ALL_A_PH | herbivory | pollination | 1.23 | 1.33 | 0.66 | |
| 5 | Sinohara_ALL_E_PH | herbivory | pollination | 1.19 | 1.06 | 0.91 | * |
| 6 | Sinohara_ALL_I_PH | herbivory | pollination | 1.20 | 0.68 | 0.95 | * |
| 7 | Sinohara_2_E_PH | pollination | herbivory | 0.98 | 0.86 | 0.76 | * |
| 8 | Sinohara_3_E_PH | pollination | herbivory | 1.35 | 0.93 | 1.13 | * |
| 9 | Sinohara_4_I_PH | herbivory | pollination | 0.77 | 0.54 | 0.67 | * |
| 10 | Melian_OO_OO_PH | pollination | herbivory | 2.08 | 1.12 | 0.75 | * |
| 11 | Hackett_1_ALL_PH | pollination | herbivory | 1.54 | 1.42 | 0.98 | * |
| 12 | Hackett_2_ALL_PH | pollination | herbivory | 0.92 | 0.83 | 0.91 | * |
| 13 | Hackett_1_S_PH | pollination | herbivory | 0.85 | 0.81 | 0.88 | |
| 14 | Hackett_1_GL_PH | pollination | herbivory | 1.10 | 1.08 | 0.58 | * |
| 15 | Pocock_OO_OO_PH | pollination | herbivory | 1.31 | 1.63 | 0.62 | |
| 16 | Melian_OO_OO_HSD | herbivory | dispersion | 1.24 | 0.75 | 1.30 | |
| 17 | McFayden_ALL_A_HPa | herbivory | parasitism | 1.29 | 0.88 | 1.39 | |
| 18 | McFayden_1_A_HPa | herbivory | parasitism | 1.16 | 0.48 | 1.24 | |
| 19 | McFayden_2_A_HPa | herbivory | parasitism | 1.21 | 0.63 | 1.35 | |
| 20 | McFayden_3_A_HPa | herbivory | parasitism | 0.98 | 0.51 | 1.10 | |
| 21 | McFayden_4_A_HPa | herbivory | parasitism | 1.01 | 0.45 | 0.98 | * |
| 22 | McFayden_5_A_HPa | herbivory | parasitism | 1.18 | 0.55 | 1.17 | * |
| 23 | McFayden_6_A_HPa | herbivory | parasitism | 1.12 | 0.50 | 1.12 | |
| 24 | McFayden_7_A_HPa | herbivory | parasitism | 1.04 | 0.44 | 0.93 | * |
| 25 | McFayden_8_A_HPa | herbivory | parasitism | 1.33 | 0.60 | 1.38 | |
| 26 | McFayden_9_A_HPa | herbivory | parasitism | 1.05 | 0.31 | 1.31 | |
| 27 | McFayden_10_A_HPa | herbivory | parasitism | 1.23 | 0.47 | 1.32 | |
| 28 | McFayden_ALL_B_HPa | herbivory | parasitism | 1.28 | 0.87 | 1.20 | * |
| 29 | McFayden_1_B_HPa | herbivory | parasitism | 0.99 | 0.47 | 1.07 | |
| 30 | McFayden_2_B_HPa | herbivory | parasitism | 1.13 | 0.51 | 0.89 | * |
| 31 | McFayden_3_B_HPa | herbivory | parasitism | 0.99 | 0.53 | 0.96 | * |
| 32 | McFayden_4_B_HPa | herbivory | parasitism | 1.16 | 0.54 | 1.05 | * |
| 33 | McFayden_5_B_HPa | herbivory | parasitism | 1.34 | 0.36 | 1.37 | |
| 34 | McFayden_6_B_HPa | herbivory | parasitism | 0.84 | 0.33 | 0.89 | |
| 35 | McFayden_7_B_HPa | herbivory | parasitism | 0.92 | 0.31 | 1.11 | |
| 36 | McFayden_8_B_HPa | herbivory | parasitism | 1.12 | 0.53 | 1.08 | * |
| 37 | McFayden_9_B_HPa | herbivory | parasitism | 0.80 | 0.51 | 0.85 | |
| 38 | McFayden_10_B_HPa | herbivory | parasitism | 0.97 | 0.34 | 1.07 | |
| 39 | Hackett_1_ALL_HPa | herbivory | parasitism | 0.64 | 0.16 | 0.73 | |
| 40 | Hackett_1_WL_HPa | herbivory | parasitism | 0.42 | 0.21 | 0.33 | * |
| 41 | Melian_OO_OO_PSD | pollination | dispersion | 1.66 | 1.12 | 1.30 | * |
| 42 | Dattilo_OO_OO_PSD | pollination | dispersion | 1.14 | 1.11 | 1.27 | |
| 43 | Dattilo_OO_OO_PA | pollination | ant | 1.09 | 1.12 | 0.79 | |

Table S2: Degree heterogeneity of the linking set species in the tripartite networks in the data-set. Different columns show the degree heterogeneity of all linking set species in the tripartite network, and also for the subset present in each of the two bipartite networks that compose the tripartite. Networks marked with a star indicates those where the degree heterogeneity of the linking set species in the merged (i.e. tripartite) network is largar than that of the subset of linking set species included in each of the two bipartite networks composing the tripartite.

| int | $\frac{\sigma_k}{<k>}$ | $Z_{NL}\left(\frac{\sigma_k}{<k>}\right)$ | r | $Z_K(r)$ | count |
|---|---|---|---|---|---|
| ant-mutualism | 0.93 | 1.18() | -4.38 | -0.26() | 1 |
| dispersion | 1.25 | 2.01(*) | -0.38 | -4.29(**) | 3 |
| herbivory | 1.14 | 2.85(*) | -0.04 | 0.19() | 41 |
| parasitism | 1.00 | 2.45(*) | 0.05 | 1.90() | 24 |
| pollination | 1.28 | 2.14(*) | -0.22 | -0.92() | 19 |

Table S3: Average degree heterogeneity $\left(\frac{\sigma_k}{<k>}\right)$ and degree-degree correlations (r) in the different interaction layers of the tripartite networks of our data set. * indicates that the the null hypothesis can be rejected with 95% confidence and ** with 98% confidence.

#### 4 Connector nodes

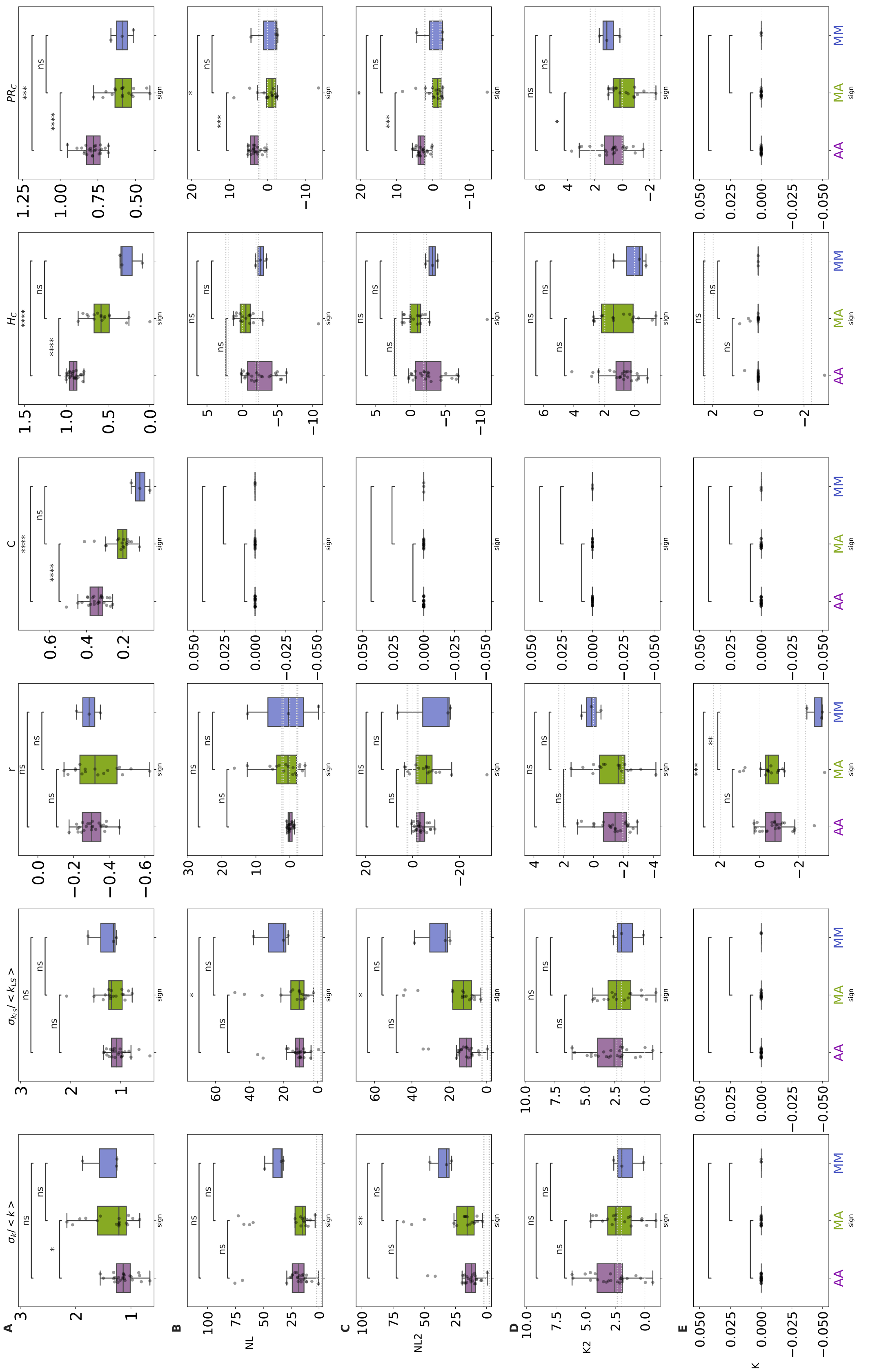


Figure S5: value and Zscore of several structural properties (columns) in the 4 different null models studied (rows). Each row is marked with a letter, A for the boxplots of the empirical values (name above each panel) and B to C the Zscore in the different null models, from the less rigid ("NL") to the most conservative ("K")

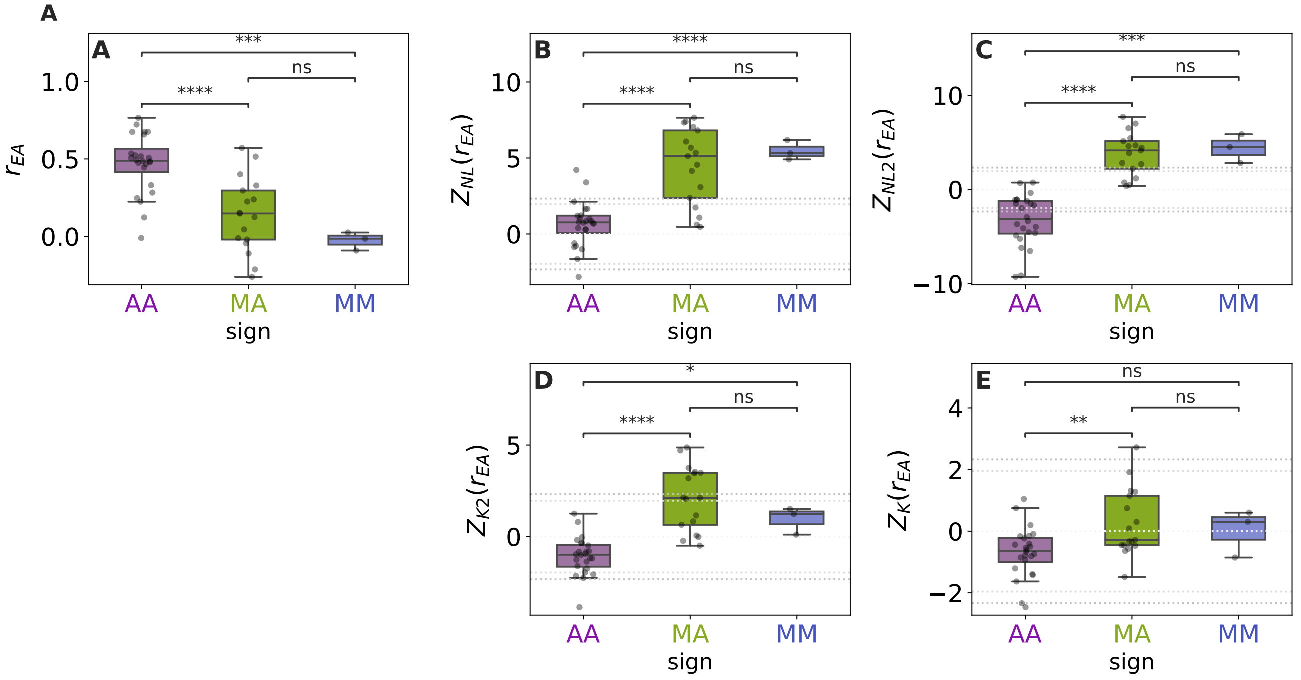


Figure S6: A: Correlation between the extinction areas of the two different interaction layers ($r_{EA}$) in the tripartite networks in our data set when plants are randomly driven to extinction, grouped and color coded by the signs of their interaction layers. B to E: Z-score of $r_{EA}$ in the different null models. The horizontal lines represent the Z-score values associated with a 95% and 98% confidence interval, z=1.96 and z=2.33 respectively.

#### 5 Interdependence

We study how the extinction areas of the two interaction layers are correlated in the tripartite networks of our database when plants were randomly driven to extinction. A positive correlation means that the same plant species that are important for the survival of animal species in one interaction layer are also important for the survival of species in the other interaction layer. However, if the relevant plants are different, deleting a given plant will cause a large loss of animal species in one layer, but not have much effect in the other layer, rendering no interdependence.

We find that, in general, the correlation between the extinction areas ($r_{EA}$) was either positive or null in all the networks. However, some MA and MM networks ($\sim$20% of all networks in the database) show a negative correlation (albeit very weak in most cases). The value of $r_{EA}$ found in AA networks (Fig. S6.A) is on average significantly higher from that found in MM and MA networks, mirroring previous results where the correlations between species sets that where linked by feeding interactions were found to be stronger [13].

Comparison with the four null models indicates that, for AA networks, the positive interdependence may be a result of the process in which secondary extinctions take place (i.e. cascading extinctions), since the empirical value is not significantly different from the null expectation in most cases (see Fig. S6 B to E). As for MM and MA networks, comparison with all null models shows that the structure of empirical networks (in particular the degree heterogeneity of the species, Fig. S6 B and C) makes the two layers more decoupled than expected by chance, since the null models predict in fact a negative correlation between the extinction areas. In the two null models where degree heterogeneity is conserved the empirical interdependence is not different from the null expectation in any case (Fig. S6 D and E).

##### Effect of structural features in interdependence

The different ways in which the two interaction layers are connected is reflected also in how the structural features of the connector nodes are able to explain the interdependence. In MM and MA networks, these structural features play an important role: $H_C$ and $PR_c$ alone account for $\sim$55% and $\sim$20% of the variance in interdependence, respectively (see Fig. S7.E and F). In AA networks, in contrast, the structural metric that best explains the variance in interdependence is $C$, but only $\sim$16% of it (see Fig. S7.D). In MM and MA networks, a multiple linear regression explains 69% of the interdependence as a function of $H_C$ (how frequently hubs are connector nodes) and $PR_c$ (the average participation ratio of the connector nodes) and disassortativity, as shown in Table S4. No structural metric explains more than 16% of the variance in interdependence in AA networks.

Regarding the presence of (weak) anti-correlations found in MA and MA networks, one could think that their origin lies in the fact that not all the targeted plant species are shared between the two layers in MA and MM networks. In those cases,

Table S4: Multiple linear regression of interdependence ($r_{EA}$) vs structural features

| | $r_{EA}$ | | | |
|---|---|---|---|---|
| | AA(all) | MA & MM(all) | AA(best) | MA & MM(best) |
| $\sigma_k / < k >$ | -0.03 | 0.19 | | |
| | (0.28) | (0.18) | | |
| $r_b$ | 0.35 | -0.32* | | -0.30** |
| | (0.24) | (0.16) | | (0.14) |
| $C$ | 0.44* | 0.12 | 0.40* | |
| | (0.22) | (0.15) | (0.20) | |
| $H_C$ | -0.42 | 0.57*** | | 0.58*** |
| | (0.25) | (0.15) | | (0.14) |
| $PR_C$ | -0.17 | 0.44** | | 0.34** |
| | (0.27) | (0.18) | | (0.13) |
| AIC | 71.21 | 39.49 | 67.92 | 38.84 |
| Observations | 24 | 20 | 24 | 20 |
| $R^2$ | 0.31 | 0.77 | 0.16 | 0.73 |
| Adjusted $R^2$ | 0.12 | 0.69 | 0.12 | 0.68 |
| Residual Std. Error | 0.96 | 0.58 | 0.96 | 0.58 |
| F Statistic | 1.62 | 9.30*** | 4.20* | 14.15*** |

*Note:* *p<0.1; **p<0.05; ***p<0.01

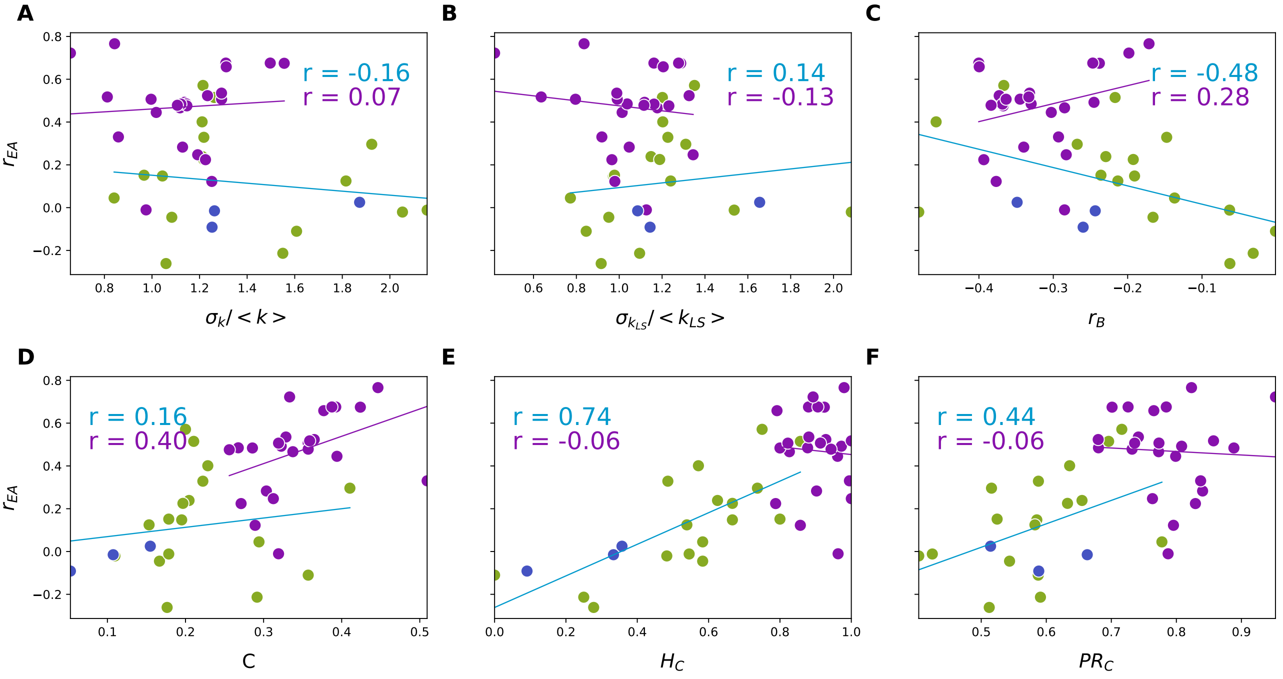


Figure S7: Correlation of interdependence ($r_{EA}$) vs the structural features studied: A) degree heterogeneity ($\sigma_k / < k >$), B) degree heterogeneity of the linking set species ($\sigma_k / < k >_{LS}$), C) degree-degree correlations ($r$), D) proportion of connector nodes inside the linking set ($C$), E) Proportion of linking set hubs that are connectors ($H_C$), and F) average participation ratio of the connector nodes ($PR_C$) on the tripartite networks in our data set. The colors of the points represent the different types of tripartite networks according to the sign of their interaction layers (see legend). The values in the lower right side are the Pearson correlation coefficients considering only MM and MA networks (blue), only AA networks (violet).

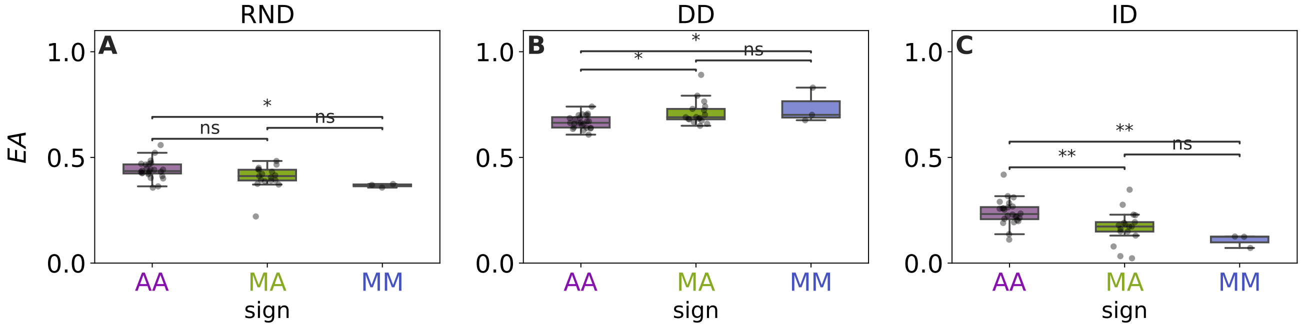


Figure S8: Extinction area of the tripartite networks in our data set in three different extinction scenarios: A) random plant extinction (RND), B) extinction of plants by decreasing degree (DD), i.e. generalist plants are attacked first, and C) extinction of plants by increasing degree (ID), i.e. specialist plants are attacked first.

secondary extinctions can happen in only one of the two interaction layers, while the other remains intact, introducing a trivial anti-correlation between the extinction areas of the two interaction layers. However, the panel D of Fig. S7 shows that there is a very low correlation between the proportion of shared species in the linking set ($C$) and the interdependence $r_{EA}$ in MA and MM networks (Pearson $r \approx 0.16$). In fact, we find that the quantity that best correlates with $r_{EA}$ in MM and MA networks ($r \approx 0.75$) is the tendency of the 20% of the most connected nodes in the linking set to be connector nodes (Fig.S7.E). That is, the fact that the generalist species in the linking set tend to be involved in the two interaction layers or not makes the difference in terms of interdependence in MA and MM networks. If the generalist plants tend to be involved in both interactions, then the interdependence is larger. If, on the contrary, the generalist plants are involved in only one of the two interaction layers, then we see no correlation, as in MM networks, or even an anti-correlation. In AA networks, on the contrary, the structural metric that best correlates with $r_{EA}$ (albeit weakly) is the proportion of connector nodes in the linking set ($C$). Since, as said before, for a parasitoid to undergo extinction the herbivore host has to go extinct first, it is not a surprise that the higher the number of connectors, the higher the interdependence of the two interaction layers.

#### 6 Robustness

All types of tripartite networks seem to react in a similar way (with an extinction area in the same range of values) to the three extinction scenarios explored, although some differences between network types were significant (see Fig. S8.A to C). In line with previous studies on networks with only one interaction type, all ecological tripartite networks were most fragile when plants were selectively attacked targeting the most connected plants first (DD scenario), and the least fragile when plants were attacked selecting the specialists plants first (ID scenario), as previously reported in mutualistic plant-pollinator, food-webs and host-parasite networks (albeit fish-parasites, not herbivore-parasites) [6, 12, 1, 3, 16]. In the random extinction scenario (RND scenario), the extinction areas showed intermediate values, with AA networks being the more fragile of the three different type of networks and MM networks the more robust (see Fig. S8.A).

To gauge the relevance of the empirical structure in determining the extinction area in the RND extinction scenario, we tested how the extinction area of the empirical networks compared to that of their different randomizations. AA empirical networks where consistently more robust than their randomizations when the original degree distribution was lost (i.e., in all null models except "K", see Fig. S9.B to D). Empirical MA and MM networks, on the other hand, were less robust than their randomizations in the "NL" ensemble (and MM networks also in the "NL2" ensemble). This indicates two things. First, that the degree distribution is a mayor driver of the extinction area, since as long as it is kept constant (null model "K") the extinction area of all types of empirical networks is not significantly different from the random expectation. Second, that the degree distribution of empirical AA networks is such that makes them more robust to the loss of plant species.

We also compared the robustness of the MA and MM tripartite networks with that of the two bipartite networks that compose the tripartite [1] to see if considering multiple interactions at the same time enhanced or hindered the robustness of the whole community. We find that, in all cases, considering the merged network did not increased the robustness to plant extinction, a result in line with previous findings in multiple-mutualistic ecological networks [4]. Rather, the robustness of the whole community ($EA_{merged}$) was in between that of the two bipartite networks composing the tripartite network (Fig. S10.D).

We investigate whether one of the two bipartite networks could be driving the values of the extinction area of the tripartite network ($EA_{merged}$), and we find that the larger (i.e. species richer) bipartite network could account for about 80% of the variance in the extinction area of the whole community (Fig. S10.A) while the extinction area of the smaller (i.e. species poorer) bipartite network could account for only 46% of the variation (Fig. S10.B). Interestingly, the robustness of the whole community could be expressed as a linear combination of the robustness of the larger and smaller bipartite networks composing the tripartite network (Fig. S10.D), with the larger one having a larger contribution to the extinction area of

[1] This analysis was only possible in MA and MM networks, since in AA networks plants appear only in one of the two bipartite networks –plant-herbivores– making it impossible to quantify the extinction area in the other bipartite network –herbivore-parasite–

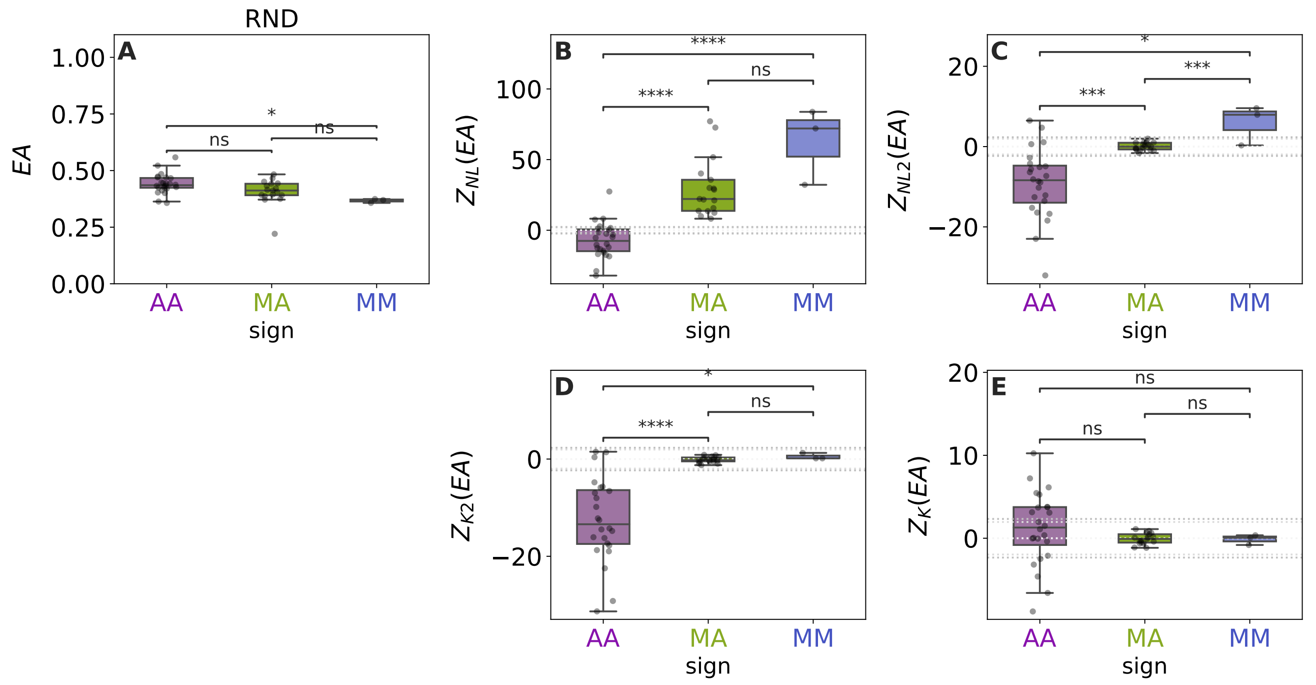


Figure S9: A: Extinction area ($EA$) of the tripartite networks in our data set when plants are randomly driven to extinction, grouped and color coded by the sign of the interaction layers. B to E: Z-score of $EA$ in the different null models. The horizontal lines represent the Z-score values associated with a 95% and 98% confidence interval, z=1.96 and z=2.33 respectively.

the tripartite network (0.74) and the smaller one a smaller contribution (0.25). As we see in Fig. S10.C, there is a very good agreement between the measured values of extinction area in the empirical tripartite networks ($EA_{merged}$) and the extinction area estimated from the two extinction areas of the ($EA_{est}$), highlighting the importance of considering the multiple interactions to better estimate the robustness of the whole community.

##### Effect of basic structural features in EA

Table S5: Multiple regression of extinction area ($EA$) vs structural features

| | $EA$ | | | |
|---|---|---|---|---|
| | AA(all) | MA & MM(all) | AA(best) | MA & MM(best) |
| $\sigma_k / < k >$ | -0.72*** | -0.60** | -0.84*** | -0.38* |
| | (0.16) | (0.28) | (0.11) | (0.19) |
| $r_b$ | -0.09 | -0.09 | | |
| | (0.14) | (0.24) | | |
| $C$ | -0.19 | 0.64** | -0.24** | 0.51** |
| | (0.13) | (0.23) | (0.11) | (0.19) |
| $H_C$ | -0.04 | -0.26 | | |
| | (0.15) | (0.22) | | |
| $PR_C$ | 0.24 | -0.25 | | |
| | (0.16) | (0.27) | | |
| AIC | 44.67 | 56.35 | 43.07 | 53.24 |
| Observations | 24 | 20 | 24 | 20 |
| $R^2$ | 0.77 | 0.46 | 0.73 | 0.38 |
| Adjusted $R^2$ | 0.71 | 0.27 | 0.70 | 0.31 |
| Residual Std. Error | 0.55 | 0.88 | 0.56 | 0.86 |
| F Statistic | 12.16*** | 2.41* | 27.77*** | 5.18** |
| *Note:* | | | *p<0.1; **p<0.05; ***p<0.01 | |

Studying how the basic structural features of the tripartite networks affect their robustness to plant loss, we find that the effects of the different structural features greatly depend on the explored extinction scenario (see Figure S11).

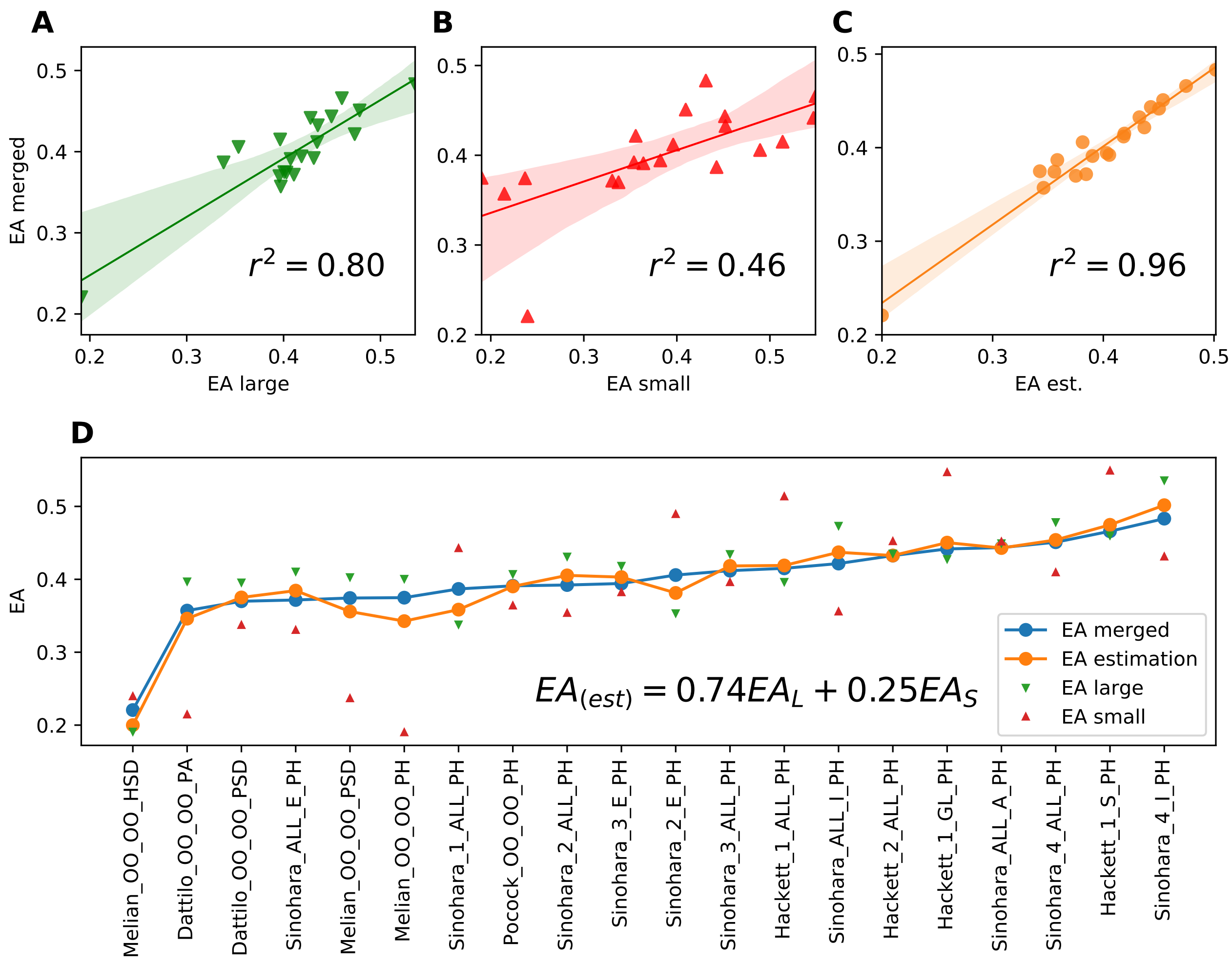


Figure S10: Upper panel: correlation of the extinction area of the whole network ($EA$ merged ) with A) the extinction area of the larger interaction layer, B) the extinction area of the smaller interaction layer, and C) with the best estimation of the extinction area as a composition of the extinction areas of the two interaction layers. Lower panel: Comparison of the extinction areas of the tripartite networks (in blue) with that of the two interaction layers composing them (EA of the larger interaction layer in green and smaller in red) and with the best estimated extinction area as a composition of the extinction areas of the two interaction layers (formula shown).

We did a multiple linear regression of the extinction area ($EA$) in the RND extinction scenario as a function of the different structural features (Table S5)

Degree heterogeneity is associated with a decrease in the fragility of all tripartite networks in the RND and ID scenarios ($r_{RND} = -0.62$, $r = -0.75$), but increasing it in MA and MM networks in the DD scenario ($r_{DD} = 0.73$). The positive effect of degree heterogeneity on robustness was already reported for bipartite mutualistic networks in [2] (through nestedness, but it was also shown that nestedness is consequence of degree heterogeneity [8]). These results are also in line with previous studies showing that ecological networks are quite robust to the extinction of the most specialist species, but quite fragile if the most generalized species were the ones going extinct [15, 6, 12]. This robustness to random extinctions has largely been explained by their heterogeneous structure, in food webs [15] and in plant-mutualistic networks [12], and is also present in the tripartite networks in our data set.

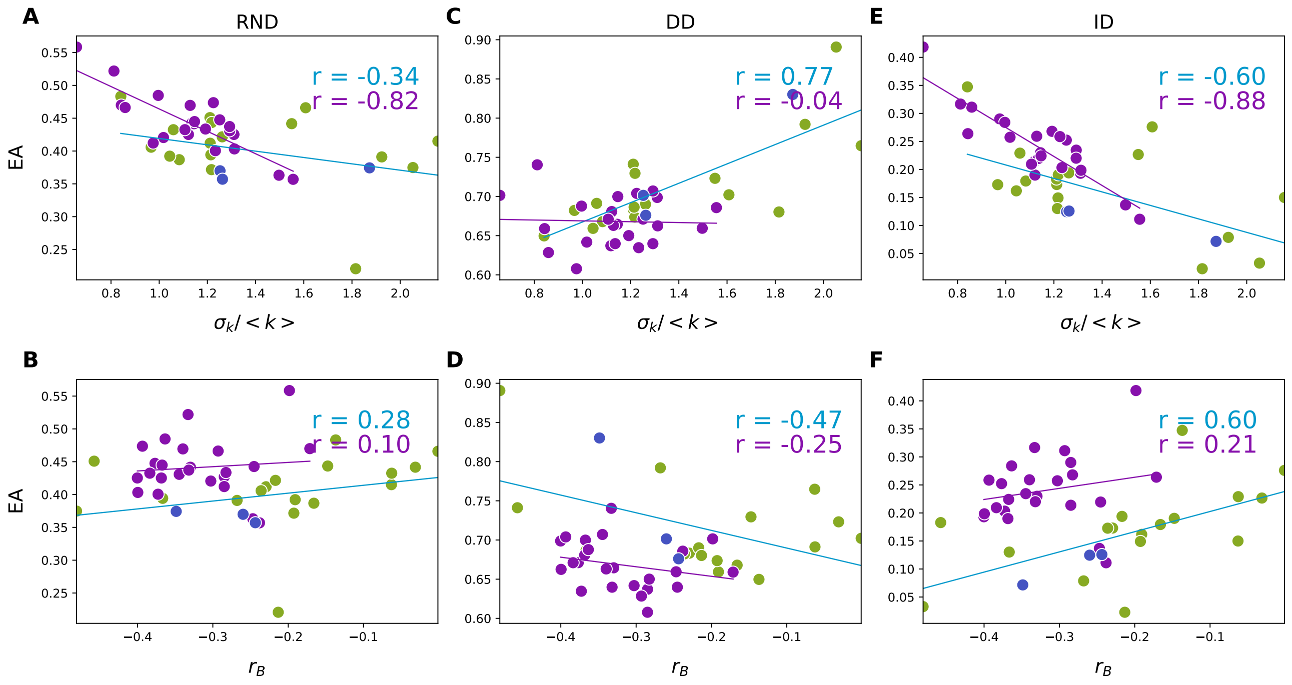


Figure S11: Effect of degree heterogeneity ($\sigma_k / < k >$) and degree-degree correlations ($r$) on the extinction area (EA) of the tripartite networks in our data set. The colors of the point represent the different types of tripartite networks according to the sign of their interaction layers (violet AA, green MA, and blue MM). The values in the upper right side are the Pearson correlation coefficients considering only MM and MA networks (blue), and only AA networks (violet). The lines represents the best linear regression for each of the correlations.

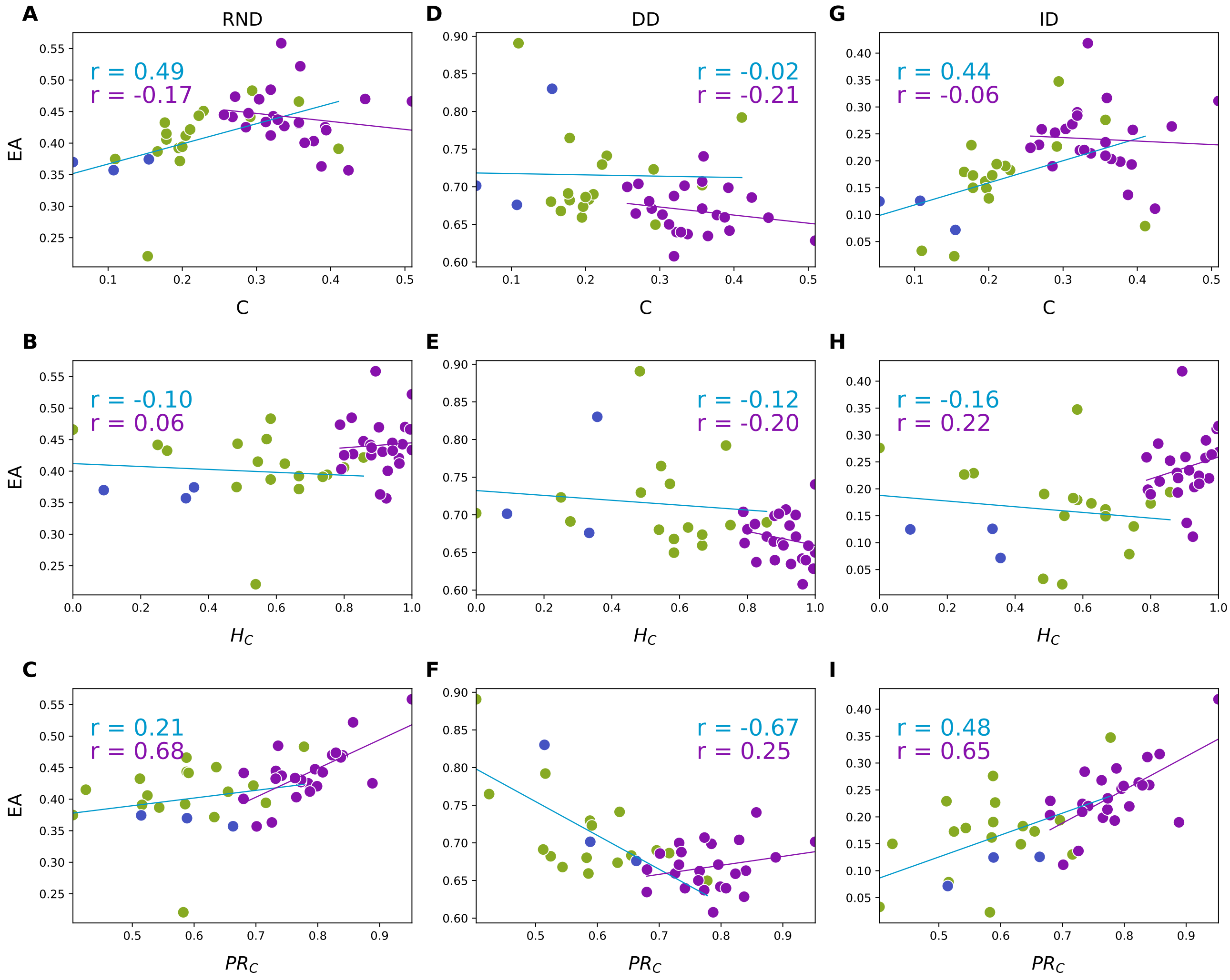


Figure S12: Effect of structural features of the linking set on the extinction area (EA) of the tripartite networks in our data set. The color of the points represents the different types of tripartite networks according to the sign of their interaction layers (violet AA, green MA, and blue MM). The values on the side are the Pearson correlation coefficients considering only MM and MA networks (blue), and only AA networks (violet). The lines represent the best linear regression for each of the correlations.

#### 7 Tables

| set | name | $\sigma_k / < k >$ | $Z$ NL |
|---|---|---|---|
| Ant | Dattilo_OO_OO_PA | 1.00 | 2.41(*) |
| Herbivore | Hackett_1_ALL_PH | 0.32 | 1.10() |
| | Hackett_1_GL_PH | 0.00 | nan() |
| | Hackett_1_S_PH | 0.26 | -0.09() |
| | Hackett_2_ALL_PH | 0.51 | 0.78() |
| | Melian_OO_OO_HSD | 1.25 | 1.36() |
| | Melian_OO_OO_PH | 1.15 | 1.31() |
| | Pocock_OO_OO_PH | 11.07 | 35.93(**) |
| | Sinohara_1_ALL_PH | 0.66 | 1.75() |
| | Sinohara_2_ALL_PH | 0.68 | 2.37(*) |
| | Sinohara_2_E_PH | 0.51 | 0.91() |
| | Sinohara_3_ALL_PH | 0.83 | 3.33(**) |
| | Sinohara_3_E_PH | 0.64 | 0.96() |
| | Sinohara_4_ALL_PH | 0.52 | 2.03(*) |
| | Sinohara_4_I_PH | 0.22 | 0.00() |
| | Sinohara_ALL_A_PH | 0.76 | 4.48(**) |
| | Sinohara_ALL_E_PH | 0.83 | 3.45(**) |
| | Sinohara_ALL_I_PH | 0.51 | 1.45() |
| Host | Hackett_1_ALL_HPa | 0.64 | 1.77() |
| | Hackett_1_WL_HPa | 0.42 | -0.42() |
| | McFayden_10_A_HPa | 1.23 | 8.25(**) |
| | McFayden_10_B_Hpa | 0.98 | 4.29(**) |
| | McFayden_1_A_HPa | 1.16 | 6.13(**) |
| | McFayden_1_B_Hpa | 0.99 | 5.32(**) |
| | McFayden_2_A_HPa | 1.21 | 5.51(**) |
| | McFayden_2_B_Hpa | 1.13 | 4.00(**) |
| | McFayden_3_A_HPa | 0.99 | 6.29(**) |
| | McFayden_3_B_Hpa | 0.99 | 3.47(**) |
| | McFayden_4_A_HPa | 1.01 | 4.50(**) |
| | McFayden_4_B_Hpa | 1.16 | 5.31(**) |
| | McFayden_5_A_HPa | 1.19 | 6.27(**) |
| | McFayden_5_B_Hpa | 1.36 | 8.06(**) |
| | McFayden_6_A_HPa | 1.12 | 5.89(**) |
| | McFayden_6_B_Hpa | 0.84 | 3.64(**) |
| | McFayden_7_A_HPa | 1.04 | 3.88(**) |
| | McFayden_7_B_Hpa | 0.92 | 6.85(**) |
| | McFayden_8_A_HPa | 1.33 | 5.62(**) |
| | McFayden_8_B_Hpa | 1.12 | 5.67(**) |
| | McFayden_9_A_HPa | 1.05 | 6.33(**) |
| | McFayden_9_B_Hpa | 0.80 | 2.29(*) |
| | McFayden_ALL_A_HPa | 1.40 | 11.78(**) |
| | McFayden_ALL_B_Hpa | 1.38 | 10.72(**) |
| Parasitoid | Hackett_1_ALL_HPa | 0.61 | 4.24(**) |
| | Hackett_1_WL_HPa | 0.33 | -0.00() |
| | McFayden_10_A_HPa | 0.55 | 1.76() |
| | McFayden_10_B_Hpa | 0.41 | 0.49() |
| | McFayden_1_A_HPa | 0.66 | 1.74() |
| | McFayden_1_B_Hpa | 0.62 | 1.75() |
| | McFayden_2_A_HPa | 0.48 | 1.29() |
| | McFayden_2_B_Hpa | 0.56 | 1.50() |
| | McFayden_3_A_HPa | 0.59 | 4.00(**) |
| | McFayden_3_B_Hpa | 0.69 | 2.72(*) |
| | McFayden_4_A_HPa | 0.68 | 3.16(**) |
| | McFayden_4_B_Hpa | 0.54 | 0.92() |
| | McFayden_5_A_HPa | 0.61 | 2.37(*) |
| | McFayden_5_B_Hpa | 0.53 | 1.53() |
| | McFayden_6_A_HPa | 0.55 | 1.86() |
| | McFayden_6_B_Hpa | 0.63 | 2.58(*) |
| | McFayden_7_A_HPa | 0.49 | 0.30() |
| | McFayden_7_B_Hpa | 0.35 | -0.98() |
| | McFayden_8_A_HPa | 0.69 | 2.83(*) |
| | McFayden_8_B_Hpa | 0.64 | 2.36(*) |
| | McFayden_9_A_HPa | 0.31 | -0.49() |
| | McFayden_9_B_Hpa | 0.48 | 2.67(*) |
| | McFayden_ALL_A_HPa | 0.98 | 7.17(**) |
| | McFayden_ALL_B_Hpa | 0.96 | 6.74(**) |
| Plant | Dattilo_OO_OO_PA | 1.08 | 3.99(**) |
| | Dattilo_OO_OO_PSD | 1.14 | 3.76(**) |
| | Hackett_1_ALL_HPa | 0.85 | 1.65() |
| | Hackett_1_ALL_PH | 1.54 | 4.78(**) |
| | Hackett_1_GL_PH | 1.10 | 2.44(*) |
| | Hackett_1_S_PH | 0.85 | 0.74() |
| | Hackett_1_WL_HPa | 0.53 | 0.26() |
| | Hackett_2_ALL_PH | 0.92 | 1.40() |
| | McFayden_10_A_HPa | 0.97 | 2.59(*) |
| | McFayden_10_B_Hpa | 1.45 | 5.43(**) |
| | McFayden_1_A_HPa | 1.46 | 4.21(**) |
| | McFayden_1_B_Hpa | 1.59 | 6.79(**) |
| | McFayden_2_A_HPa | 1.45 | 6.21(**) |
| | McFayden_2_B_Hpa | 0.74 | 2.62(*) |
| | McFayden_3_A_HPa | 1.41 | 5.43(**) |
| | McFayden_3_B_Hpa | 1.74 | 8.52(**) |
| | McFayden_4_A_HPa | 1.08 | 4.64(**) |
| | McFayden_4_B_Hpa | 1.13 | 3.85(**) |
| | McFayden_5_A_HPa | 1.14 | 5.50(**) |
| | McFayden_5_B_Hpa | 1.11 | 3.88(**) |
| | McFayden_6_A_HPa | 1.20 | 5.06(**) |
| | McFayden_6_B_Hpa | 0.92 | 3.44(**) |
| | McFayden_7_A_HPa | 1.39 | 5.50(**) |
| | McFayden_7_B_Hpa | 0.81 | 2.48(*) |
| | McFayden_8_A_HPa | 1.09 | 3.11(**) |
| | McFayden_8_B_Hpa | 1.14 | 3.76(**) |
| | McFayden_9_A_HPa | 1.16 | 4.34(**) |
| | McFayden_9_B_Hpa | 1.08 | 3.23(**) |
| | McFayden_ALL_A_HPa | 1.94 | 12.74(**) |
| | McFayden_ALL_B_Hpa | 1.80 | 11.38(**) |
| | Melian_OO_OO_HSD | 1.24 | 8.70(**) |
| | Melian_OO_OO_PH | 2.08 | 9.12(**) |
| | Melian_OO_OO_PSD | 1.66 | 5.03(**) |
| | Pocock_OO_OO_PH | 1.31 | 2.71(*) |
| | Sinohara_1_ALL_PH | 0.95 | 2.41(*) |
| | Sinohara_2_ALL_PH | 0.97 | 2.48(*) |
| | Sinohara_2_E_PH | 0.98 | 1.83() |
| | Sinohara_3_ALL_PH | 1.15 | 2.91(*) |
| | Sinohara_3_E_PH | 1.35 | 4.15(**) |
| | Sinohara_4_ALL_PH | 1.20 | 3.22(**) |
| | Sinohara_4_I_PH | 0.77 | 1.01() |
| | Sinohara_ALL_A_HPa | 1.23 | 3.15(**) |
| | Sinohara_ALL_E_PH | 1.19 | 3.48(**) |
| | Sinohara_ALL_I_PH | 1.20 | 2.95(*) |
| Pollinator | Dattilo_OO_OO_PA | 1.28 | 11.46(**) |
| | Dattilo_OO_OO_PSD | 1.28 | 12.44(**) |
| | Hackett_1_ALL_PH | 1.12 | 12.91(**) |
| | Hackett_1_GL_PH | 0.78 | 4.57(**) |
| | Hackett_1_S_PH | 0.67 | 4.08(**) |
| | Hackett_2_ALL_PH | 0.93 | 5.83(**) |
| | Melian_OO_OO_PH | 1.04 | 10.83(**) |
| | Melian_OO_OO_PSD | 1.04 | 10.39(**) |
| | Pocock_OO_OO_PH | 1.16 | 13.93(**) |
| | Sinohara_1_ALL_PH | 1.23 | 4.21(**) |
| | Sinohara_2_ALL_PH | 1.08 | 3.73(**) |
| | Sinohara_2_E_PH | 0.88 | 2.40(*) |
| | Sinohara_3_ALL_PH | 1.28 | 6.31(**) |
| | Sinohara_3_E_PH | 1.12 | 4.37(**) |
| | Sinohara_4_ALL_PH | 0.92 | 2.50(*) |
| | Sinohara_4_I_PH | 0.81 | 1.37() |
| | Sinohara_ALL_A_HPa | 0.80 | 2.97(*) |
| | Sinohara_ALL_E_PH | 1.28 | 4.00(**) |
| | Sinohara_ALL_I_PH | 1.07 | 3.31(**) |
| Seed disperser | Dattilo_OO_OO_PSD | 0.70 | 1.80() |
| | Melian_OO_OO_HSD | 0.78 | 1.36() |
| | Melian_OO_OO_PSD | 1.13 | 2.32(*) |

Table S6: Degree Heterogeneity and degree-degree correlations in the tripartite networks of our database.

| int | name | $\sigma_k/ < k >$ | $Z(\sigma_k/ < k >)$ NL | r | $Z(r)$ NL | $Z(r)$ K |
|---|---|---|---|---|---|---|
| ant-mutualism | Dattilo_OO_OO_PA | 0.93 | 1.18() | -0.33 | -2.74(*) | -0.24() |
| dispersion | Dattilo_OO_OO_PSD | 1.15 | 1.78() | -0.34 | -0.63() | -0.67() |
| | Melian_OO_OO_HSD | 1.30 | 2.39(*) | -0.52 | -1.41() | -5.50(**) |
| | Melian_OO_OO_PSD | 1.30 | 1.87() | -0.52 | -1.34() | -5.86(**) |
| herbivory | Hackett_1_ALL_HPa | 0.93 | 1.47() | -0.47 | 1.85() | -0.44() |
| | Hackett_1_ALL_PH | 1.06 | 1.30() | -0.34 | 2.90(*) | 2.77(*) |
| | Hackett_1_GL_PH | 0.55 | 0.41() | -0.39 | 1.07() | inf(**) |
| | Hackett_1_S_PH | 0.98 | 0.65() | -0.46 | 2.00() | 1.74() |
| | Hackett_1_WL_HPa | 0.78 | 0.29() | -0.65 | 0.36() | -0.89() |
| | Hackett_2_ALL_PH | 0.78 | 1.46() | 0.06 | 0.34() | 3.02(**) |
| | McFayden_10_A_HPa | 1.09 | 2.11(*) | -0.49 | 0.87() | -1.48() |
| | McFayden_10_B_Hpa | 1.40 | 5.38(**) | -0.32 | 3.08(**) | 1.67() |
| | McFayden_1_A_HPa | 1.61 | 3.93(**) | -0.44 | 3.06(**) | 0.01() |
| | McFayden_1_B_Hpa | 1.58 | 5.43(**) | -0.33 | 3.09(**) | 2.50(*) |
| | McFayden_2_A_HPa | 1.34 | 4.59(**) | -0.39 | -0.73() | -1.18() |
| | McFayden_2_B_Hpa | 0.70 | 1.64() | -0.25 | -1.36() | -0.54() |
| | McFayden_3_A_HPa | 1.42 | 4.46(**) | -0.32 | 3.44(**) | -0.33() |
| | McFayden_3_B_Hpa | 1.57 | 6.60(**) | -0.26 | 1.45() | 0.50() |
| | McFayden_4_A_HPa | 0.97 | 3.08(**) | -0.36 | -1.66() | -1.99() |
| | McFayden_4_B_Hpa | 1.08 | 3.07(**) | -0.38 | -0.24() | -2.59(*) |
| | McFayden_5_A_HPa | 1.08 | 3.85(**) | -0.34 | 0.14() | -1.87() |
| | McFayden_5_B_Hpa | 1.02 | 2.97(*) | -0.30 | 1.09() | -1.11() |
| | McFayden_6_A_HPa | 1.12 | 4.17(**) | -0.26 | 1.54() | -0.49() |
| | McFayden_6_B_Hpa | 0.86 | 2.53(*) | -0.25 | 2.07(*) | 0.17() |
| | McFayden_7_A_HPa | 1.28 | 5.09(**) | -0.27 | 1.84() | 0.09() |
| | McFayden_7_B_Hpa | 0.75 | 1.84() | -0.29 | 1.17() | -0.26() |
| | McFayden_8_A_HPa | 1.09 | 2.75(*) | -0.37 | 0.24() | 0.40() |
| | McFayden_8_B_Hpa | 1.12 | 2.85(*) | -0.41 | 0.19() | -0.60() |
| | McFayden_9_A_HPa | 1.17 | 3.10(**) | -0.40 | 1.83() | -0.04() |
| | McFayden_9_B_Hpa | 1.07 | 2.83(*) | -0.31 | 1.75() | -0.10() |
| | McFayden_ALL_A_HPa | 1.94 | 6.68(**) | -0.30 | 5.70(**) | -0.31() |
| | McFayden_ALL_B_Hpa | 1.83 | 6.24(**) | -0.27 | 6.02(**) | 0.58() |
| | Melian_OO_OO_HSD | 2.31 | 1.38() | -0.65 | 7.65(**) | 2.33(*) |
| | Melian_OO_OO_PH | 2.31 | 1.15() | -0.65 | 7.30(**) | 2.31(*) |
| | Pocock_OO_OO_PH | 1.34 | 3.00(**) | -0.57 | -10.57(**) | 3.09(**) |
| | Sinohara_1_ALL_PH | 0.86 | 2.49(*) | -0.25 | -3.46(**) | -1.14() |
| | Sinohara_2_ALL_PH | 0.71 | 0.94() | -0.19 | -0.04() | -0.02() |
| | Sinohara_2_E_PH | 0.68 | 1.14() | -0.22 | -1.19() | -0.28() |
| | Sinohara_3_ALL_PH | 0.97 | 2.92(*) | -0.17 | -0.82() | 0.59() |
| | Sinohara_3_E_PH | 0.91 | 2.27(*) | -0.17 | -2.27(*) | 0.88() |
| | Sinohara_4_ALL_PH | 1.00 | 2.72(*) | -0.32 | -0.00() | -1.88() |
| | Sinohara_4_I_PH | 0.53 | 0.27() | -0.35 | 0.49() | 0.49() |
| | Sinohara_ALL_A_PH | 1.22 | 3.82(**) | -0.26 | -0.27() | 0.10() |
| | Sinohara_ALL_E_PH | 0.97 | 3.00(*) | -0.07 | -1.55() | 1.19() |
| | Sinohara_ALL_I_PH | 0.71 | 0.94() | -0.19 | 1.12() | 0.82() |
| parasitism | Hackett_1_ALL_HPa | 0.68 | 1.66() | -0.26 | -2.39(*) | -0.89() |
| | Hackett_1_WL_HPa | 0.32 | 0.31() | -0.33 | -inf(**) | -0.93() |
| | McFayden_10_A_Hpa | 1.24 | 3.00(**) | -0.24 | 1.40() | 3.25(**) |
| | McFayden_10_B_Hpa | 0.96 | 1.96() | -0.30 | 0.26() | 1.73() |
| | McFayden_1_A_Hpa | 1.07 | 2.63(*) | -0.16 | -0.84() | 3.00(*) |
| | McFayden_1_B_Hpa | 0.95 | 2.60(*) | -0.13 | -0.35() | 2.76(*) |
| | McFayden_2_A_Hpa | 1.23 | 3.14(**) | -0.23 | 1.55() | 2.18(*) |
| | McFayden_2_B_Hpa | 0.90 | 1.07() | -0.25 | 1.44() | 1.26() |
| | McFayden_3_A_Hpa | 1.01 | 3.08(**) | -0.17 | 0.13() | 1.53() |
| | McFayden_3_B_Hpa | 0.92 | 1.80() | -0.15 | 0.43() | 1.92() |
| | McFayden_4_A_Hpa | 0.95 | 2.16(*) | -0.09 | 1.69() | 2.96(*) |
| | McFayden_4_B_Hpa | 0.95 | 1.68() | -0.20 | 0.33() | 2.21(*) |

| int | name | $\sigma_k / < k >$ | $Z(\sigma_k / < k >)$ NL | r | $Z(r)$ NL | $Z(r)$ K |
|---|---|---|---|---|---|---|
| | McFayden_5_A_Hpa | 1.05 | 2.98(*) | -0.20 | -0.08() | 1.59() |
| | McFayden_5_B_Hpa | 1.29 | 3.08(**) | -0.28 | 0.94() | 1.10() |
| | McFayden_6_A_Hpa | 1.02 | 2.20(*) | -0.28 | -0.03() | 0.44() |
| | McFayden_6_B_Hpa | 0.81 | 2.01(*) | -0.29 | -2.39(*) | -0.99() |
| | McFayden_7_A_Hpa | 0.86 | 1.12() | -0.28 | -0.26() | 1.28() |
| | McFayden_7_B_Hpa | 0.91 | 2.83(*) | -0.16 | -1.07() | 0.75() |
| | McFayden_8_A_Hpa | 1.33 | 2.95(*) | -0.26 | 2.04(*) | 4.19(**) |
| | McFayden_8_B_Hpa | 1.02 | 2.07(*) | -0.13 | 1.64() | 3.67(**) |
| | McFayden_9_A_Hpa | 1.12 | 3.47(**) | -0.22 | 1.48() | 1.61() |
| | McFayden_9_B_Hpa | 0.80 | 1.27() | -0.09 | 2.07(*) | 2.75(*) |
| | McFayden_ALL_A_Hpa | 1.38 | 5.49(**) | -0.11 | -1.93() | 2.14(*) |
| | McFayden_ALL_B_Hpa | 1.26 | 4.24(**) | -0.09 | 0.38() | 2.65(*) |
| pollination | Dattilo_OO_OO_PA | 1.27 | 3.57(**) | -0.26 | 1.92() | -2.74(*) |
| | Dattilo_OO_OO_PSD | 1.27 | 3.53(**) | -0.26 | 1.91() | -2.76(*) |
| | Hackett_1_ALL_PH | 2.05 | 2.48(*) | -0.41 | 13.77(**) | -0.52() |
| | Hackett_1_GL_PH | 1.52 | 1.93() | -0.50 | 5.13(**) | -0.87() |
| | Hackett_1_S_PH | 1.51 | 1.01() | -0.59 | 5.03(**) | 0.85() |
| | Hackett_2_ALL_PH | 1.06 | 1.58() | -0.41 | 2.02(*) | -0.52() |
| | Melian_OO_OO_PH | 1.93 | 1.16() | -0.37 | 16.31(**) | -1.75() |
| | Melian_OO_OO_PSD | 1.93 | 1.54() | -0.37 | 15.75(**) | -1.84() |
| | Pocock_OO_OO_PH | 2.20 | 3.32(**) | -0.51 | 8.91(**) | -1.96() |
| | Sinohara_1_ALL_PH | 1.03 | 2.45(*) | -0.22 | -2.36(*) | 1.33() |
| | Sinohara_2_ALL_PH | 1.00 | 2.53(*) | -0.28 | -3.30(**) | 0.06() |
| | Sinohara_2_E_PH | 0.86 | 1.90() | -0.31 | -3.28(**) | -1.02() |
| | Sinohara_3_ALL_PH | 1.05 | 2.28(*) | -0.27 | -2.00() | -0.95() |
| | Sinohara_3_E_PH | 1.02 | 2.71(*) | -0.39 | -2.91(*) | -1.35() |
| | Sinohara_4_ALL_PH | 1.00 | 1.88() | -0.45 | -3.49(**) | -1.75() |
| | Sinohara_4_I_PH | 0.72 | 1.08() | -0.23 | -2.05(*) | -0.05() |
| | Sinohara_ALL_A_PH | 0.76 | 0.87() | -0.10 | 0.36() | 0.65() |
| | Sinohara_ALL_E_PH | 1.08 | 2.76(*) | -0.26 | -3.84(**) | -0.64() |
| | Sinohara_ALL_I_PH | 1.01 | 2.11(*) | -0.36 | -3.35(**) | -0.95() |

Table S7: Degree heterogeneity and degree-degree correlations in the different interaction layers of the tripartite networks of our database.
